# Apple peel and flesh contain pro-neurogenic compounds

**DOI:** 10.1101/2020.06.25.170266

**Authors:** Muhammad Ichwan, Tara L. Walker, Zeina Nicola, Jutta Ludwig-Müller, Christoph Böttcher, Rupert W. Overall, Vijay S Adusumilli, Merve Bulut, Enrique A. Lugo-Hernández, Gerardo Ramirez-Rodriguez, Gerd Kempermann

## Abstract

As mammals evolved exposed to particular diets, naturally abundant compounds may have become part of the set of environmental co-determinants that shaped brain structure and function. Here we investigated whether bioactive factors found in apples directly affect hippocampal neural stem cells and promote neurogenesis in the adult mouse. Whereas the consumption of apple juice *per se* neither altered adult hippocampal neurogenesis nor improved learning and memory, we did find specific direct effects of apple-derived factors on neural stem cell survival and differentiation. Our results revealed that quercetin, the most abundant flavanol in apple peel, was anti-proliferative at high concentrations but acted pro-neurogenically at low concentrations. This was confirmed *in vivo*, with intraperitoneally-delivered quercetin promoting survival and neuronal differentiation, without affecting proliferation, likely via the PI3 kinase-Akt and Nrf2-Keap1 pathways, respectively. Using a bio-assay-guided fractionation approach with high-resolution collision induced dissociation mass spectroscopy, we also identified additional pro-neurogenic compounds in apple flesh that were not related to flavonoids. In particular, we found that 3,5-dihydroxybenzoic acid, a weak agonist to the lactate receptor, significantly increased both *in vitro* and *in vivo* neural precursor cell proliferation and neurogenesis. Altogether, this work shows that both flavonoids and 3,5-dihydroxybenzoic acid are pro-neurogenic, not only by activating precursor cell proliferation but also through promoting cell cycle exit, cellular survival, and neuronal differentiation.

## Introduction

“An apple a day keeps the doctor away” – there may be some truth to this aphorism. Apart from being a source of energy, food is also believed to influence an individual’s physical and intellectual fitness. A growing number of studies have demonstrated the health benefits of phytochemicals, the chemical substances found in plants, and have attributed this to their role as substrates for biochemical reactions, cofactors or inhibitors of enzymatic reactions, ligands of membrane or intracellular receptors, scavengers of reactive chemicals, and modulators of gut microbes (for review see (Dillard & German, 2000).

Active dietary compounds are also vital for maintaining cognitive function. One of the processes underlying this maintenance is brain plasticity, by which structural and functional modifications occur in response to internal and external stimuli. Adult hippocampal neurogenesis is a particular form of brain plasticity whereby functional neurons are generated throughout life and integrated into the existing neuronal circuitry. This process drives cellular and synaptic plasticity in the hippocampus and plays a role in mediating particular forms of learning and memory. Flavonoids, the abundant phytonutrients found in fruits and vegetables, can modulate molecular signaling pathways which influence these cognitive processes (Williams et al., 2004; Spencer, 2009). Prominent examples of flavonoids and polyphenols are resveratrol (found in the skin of red grapes and consequently in red wine) and epigallo-catechin-3-gallate (EGCG; found in green tea). In the case of resveratrol, the biological activity has been traced to the intracellular regulation of Sirt1 and other sirtuins (Prozorovski et al., 2008). We and others have shown that EGCG and resveratrol affect adult hippocampal neurogenesis (Ortiz-Lopez et al., 2016; Torres-Perez et al., 2015; Yoo et al., 2010). However, although some positive indications have been reported (van der Made, Plat, & Mensink, 2015). One reason, besides questions of bioavailability, might be that plants actually contain a very large number of potentially active compounds. Georgiev et al., for example, lists 20 flavonoids from grapes alone (Georgiev, Ananga, & Tsolova, 2014), that likely work in concert to exert a positive effect.

As apples are one of the most widely consumed fruit worldwide, resulting in a generalized exposure across cultures, we here investigated whether apples contain substances that might sustain or promote adult hippocampal neurogenesis. We took a two-pronged approach, first investigating quercetin, the most prominent flavonoid present in apples, and secondly, extending our investigation to identify additional pro-neurogenic factors that might be present in the fruit. Specifically, and because flavonoids are largely confined to the apple peel, we investigated whether pro-neurogenic compounds can also be found in the flesh.

## Results

### Quercetin promotes cell cycle exit and promotes neuronal differentiation in vitro

Quercetin is the most abundant flavonoid in apple peel (Bhagwat 2011) and has already been linked to several neuroprotective mechanisms, particularly in relation to microglial function (Nichols et al., 2015). This class of phytochemicals can also exert an antioxidant effect due to its capacity to scavenge reactive oxygen species. Although quercetin and other natural compounds might affect adult neurogenesis through more than one pathway, we focused on their effect on stem cells, from which the adult-born neurons in the hippocampus originate. We first used the monolayer culture system, which consists of a relatively pure population of putative stem and progenitor cells to establish a dose response curve (Babu, Cheung, Kettenmann, Palmer, & Kempermann, 2007). In line with reports of inhibitory effects of high doses of quercetin on cancer stem cells, we found that incorporation of the thymidine analogue 5-bromo-2’-deoxyuridine (BrdU), as a measure of proliferative activity, was reduced at concentrations of 25 μM and higher (0 μM quercetin 52.23 ±2.42 vs 25 μM quercetin 33.53 ± 2.12 % BrdU^+^ cells, n = 4 experiments, *p* < 0.05; Fig 1A, B). At concentrations up to 35 μM, this was not due to an increased cytotoxicity, which however was detected at even higher doses with the lactate dehydrogenase assay (0 μM quercetin 13.46 ± 1.3 vs 50 μM quercetin 49.59 ± 9.8 %, n = 4, *p* < 0.01; Fig 1C) and 7-AAD staining (Fig 1D). Cell cycle analysis based on propidium iodide incorporation and flow cytometric analysis revealed that 25μM quercetin reduced S-phase entry (0 μM quercetin 14.34 ± 0.28 vs 25 μM quercetin 7.53 ± 0.3 %, n = 3 experiments, *p* < 0.001, Fig 1E) and instead caused the cells to accumulate in the G1 phase (0 μM quercetin 67.85 ± 0.66 vs 25 μM quercetin 72.31 ± 0.54 %, n = 3 experiments, *p* < 0.001, Fig 1E). Western blot analysis for cell cycle-related proteins showed that CDK4 levels were decreased and cyclinD1 remained unchanged (Fig 1F). In contrast, the cell cycle inhibitor p27 was upregulated by 25 μM quercetin, to an even greater extent than by growth factor withdrawal which causes to cells to exit the cell cycle and differentiate.

**Figure 1.**
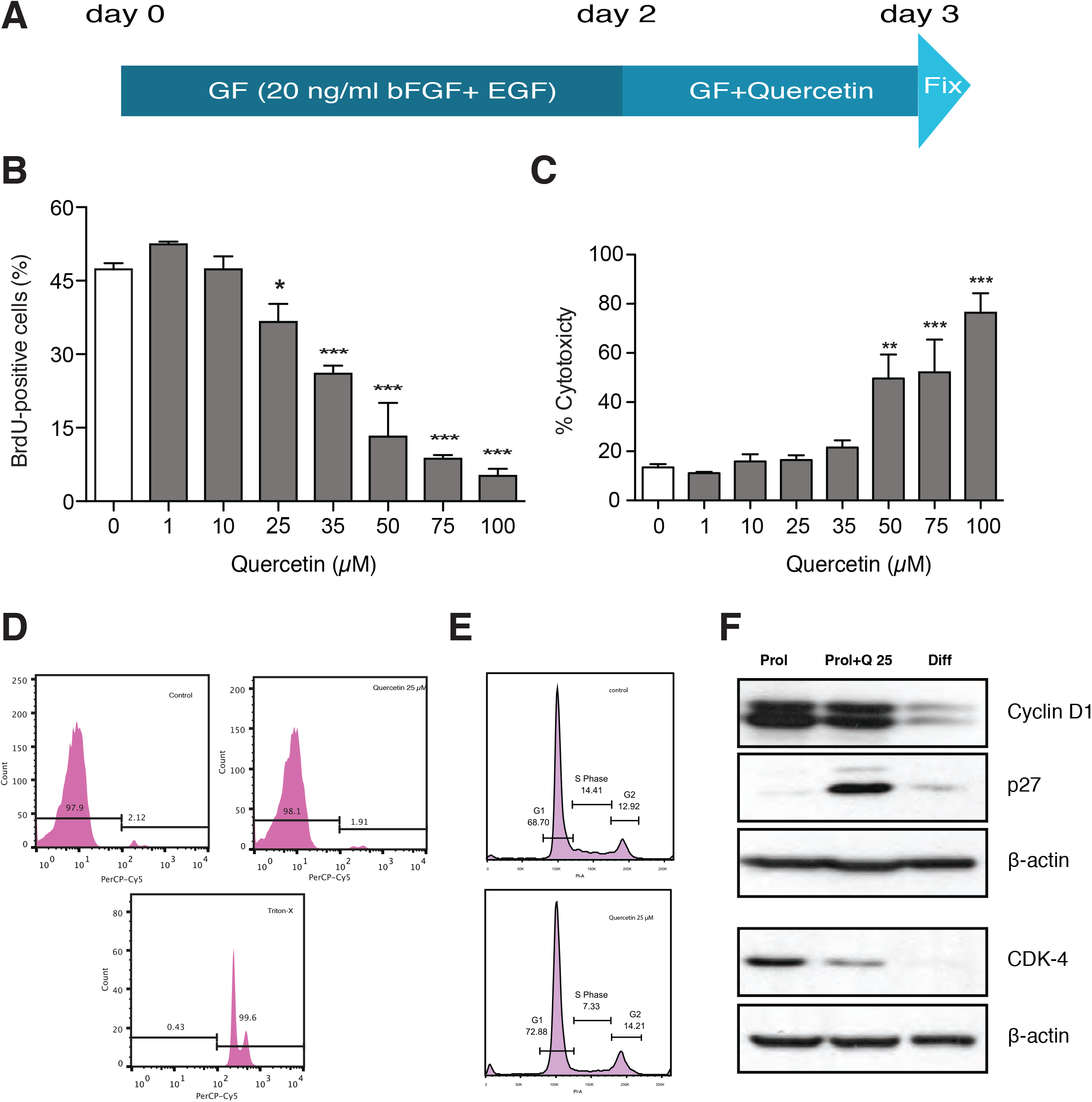
Quercetin inhibits the cell cycle without inducing cell death. (A) Schematic of the monolayer neural precursor cell culture with 24 h quercetin treatment. (B) Quercetin decreased BrdU-labeled cells dose dependently (C) Quercetin concentrations of ≥50 μM significantly increased the number of dead cells measured using the LDH cytotoxicity assay. Error bars represent SEM. (D) Cell cycle analysis using propidium iodide staining revealed a decreased number of cells entering S phase after 24 h of 25 μM quercetin treatment. (E) 7-AAD staining confirmed that quercetin (25 μM) does not induce cell death. (F) Western blot analysis of cell cycle-related proteins revealed that quercetin treatment resulted in the downregulation of CDK4, and the upregulation of p27. Cells cultured for 24 h without growth factors (Diff) were used as a control for cell cycle arrest. * *p* < 0.05, ** *p* < 0.01, *** *p* < 0.001; one-way ANOVA with Dunnett’s post-hoc test. GF = growth factors.

Together the above data indicate that quercetin upregulates the cell cycle ‘brake’ signal to counteract the mitogenic effects of growth factors. This was further supported by our observation that, in the presence of growth factors, 25 μM quercetin increased the number of cells with neuron-like morphology (n = 5 experiments, *p* < 0.001, Fig 2A, B) and βIII-tubulin expression (Fig 2C; an immature neuronal marker), whereas expression of the *bona-fide* stem cell-marker nestin was decreased (n = 6 experiments, *p* < 0.001, Fig 2D). Expression of the astrocytic marker GFAP also increased. Taken together, these findings indicate that quercetin can act pro-neurogenically even in the presence of growth factors, which normally maintain *in vitro* precursor cells in an undifferentiated and proliferative state.

**Figure 2.**
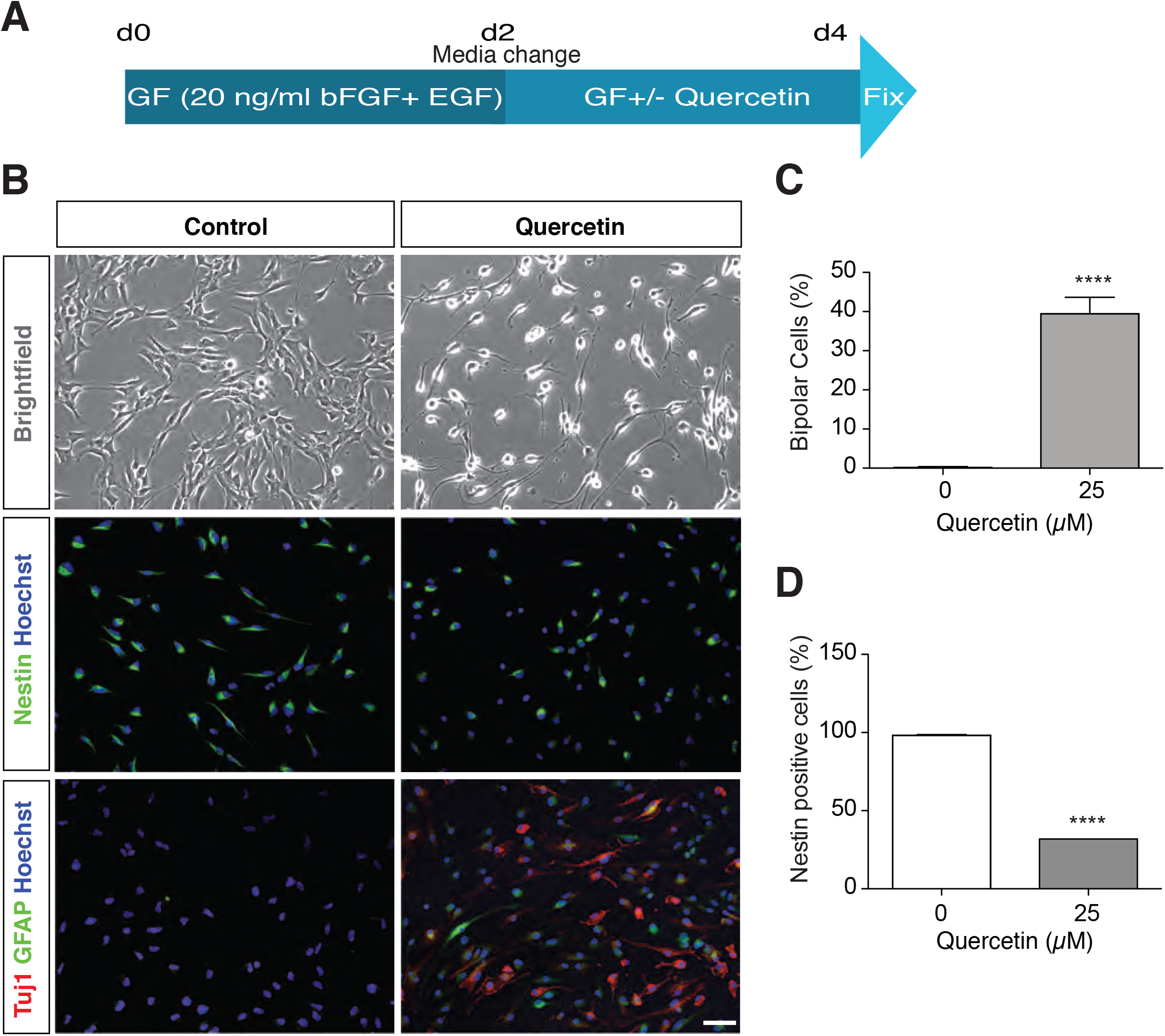
Quercetin induces neural precursor cell differentiation. (A) Experimental scheme. (B) Neural precursor cells change their morphological characteristics following quercetin treatment into cells with more “neuron-like" bipolar characteristics, (upper panels). In addition, they downregulate of nestin expression (middle panels) and upregulate expression of the neuronal marker β-III-tubulin (Tuj1) and the astrocyte marker GFAP (lower panels). Scale bar 50 μm. (C) Quercetin (25 μM) significantly increased the percentage of cells with a bipolar morphology. (D) Quercetin (25 μM) significantly decreased the percentage of Nestin^+^ cells. Error bars represent SEM. *\*\*\* p* < 0.001; Student’s t-test.

### Quercetin promotes in vitro cell survival and stimulates adult neurogenesis in vivo

The neural precursor cells in monolayer cultures can be induced to differentiate into neurons or astrocytes by removal of growth factors from the medium. However, growth factor withdrawal is a stressful condition that leads to significant cell death. As polyphenols have been shown to exert neuroprotective effects, we next examined whether quercetin supports cell survival *in vitro* during differentiation. We found that the addition of 25 μM quercetin for 4 days following growth factor withdrawal almost doubled the percentage of surviving cells (Fig 3A-C, n = 5 experiments, *p*<0.001).

**Figure 3.**
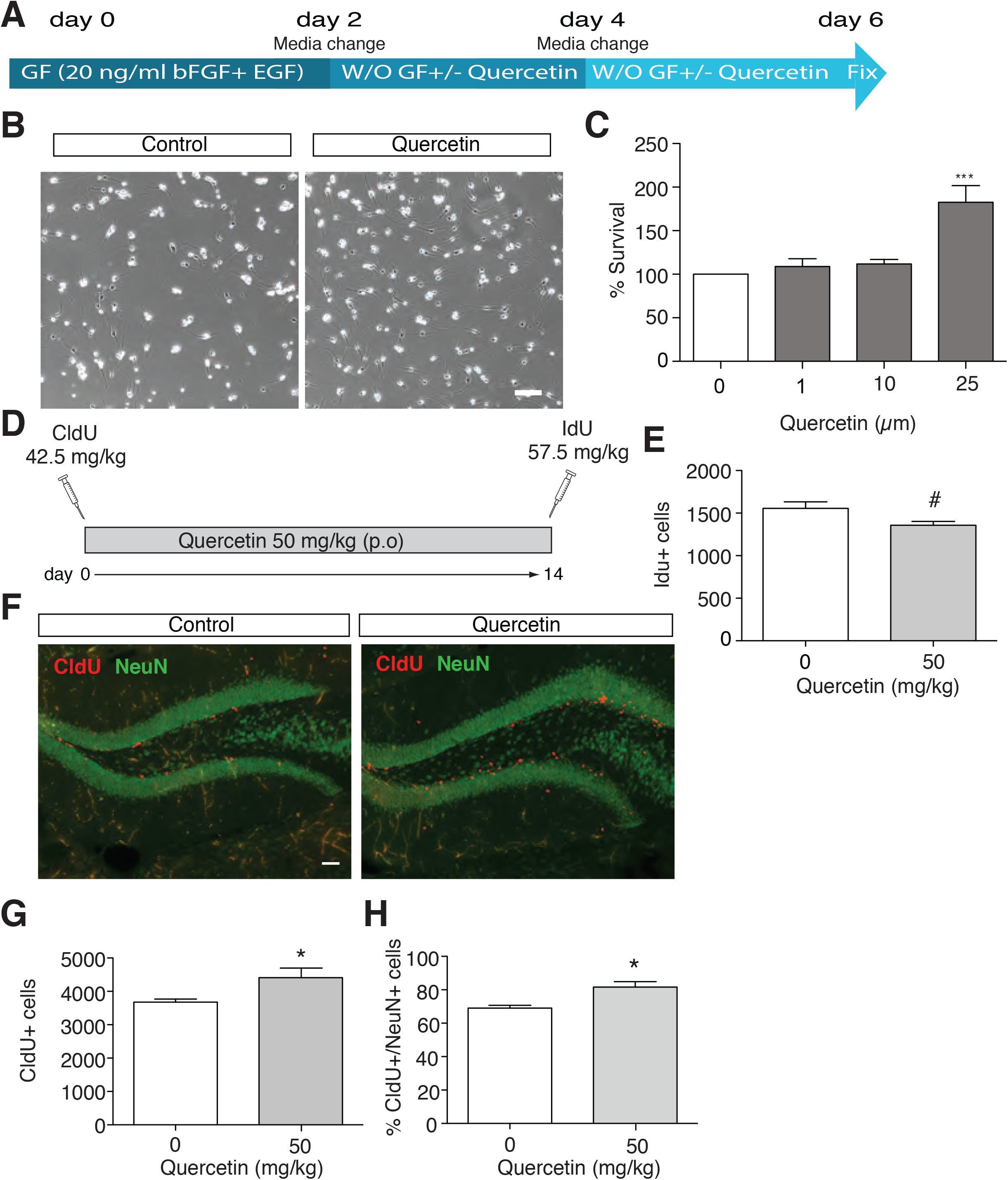
Quercetin promotes *in vitro* cell survival and stimulates adult neurogenesis *in vivo*. (A) Experimental scheme of monolayer neural precursor cells cultured under differentiation conditions. (B) Brightfield images of neural precursor cells 96 h after growth factor withdrawal. (C) Quercetin (25 μM) increased cell survival during differentiation. Bar graph shows the percentage of Resazurin fluorescence intensity compared to control. *** *p* < 0.001; one-way ANOVA with Dunnett’s post-hoc test. (D) Experimental scheme for CldU and IdU injection and quercetin treatment. (E) A trend towards a decrease in IdU+ cells was observed in the quercetin-treated animals (F) Immunofluorescence staining shows the cohort of cells that were CldU^+^ (red) and NeuN^+^ (green). A significant increase in the number of CldU^+^ (G) and CldU^+^NeuN^+^ (H) cells was observed in the quercetin-treated group. Error bars represent SEM. * *p* < 0.05, # *p* = 0.068; Student’s t-test. Scale bars are 50 μm.

Given the positive effects of quercetin on cell survival and neurogenesis *in vitro*, we next examined its effects *in vivo*. Quercetin was administered orally at 50 mg/kg, a dose previously shown to lead to sufficient accumulation of metabolites in the brain (Ishisaka et al., 2011). Proliferating cells were labeled with IdU after 14 days of quercetin treatment, whereas the effect of the treatment on cell survival was assessed in a cohort of cells labeled with CldU immediately before the treatment began (Fig 3D). After two weeks of quercetin treatment we observed a trend towards a decrease in proliferation (0 μM quercetin 1556 ± 76.77 vs 50 μM quercetin 1358 ± 45.52 IdU^+^ cells, n = 4 mice per group, *p* = 0.068; Fig 3E). In contrast, the number of CldU^+^ cells was significantly increased after quercetin treatment (0 μM quercetin 3672 ± 94.58 vs 50 μM quercetin 4410 ± 286.1 CldU^+^ cells, n = 4 mice per group, *p* = 0.049; Fig 3F). Quercetin treatment resulted in a net increase in the number of newly born neurons, with a significantly higher percentage of NeuN^+^/CldU^+^ cells compared to the control group (0 μM quercetin 69 ± 1.73 vs 50 μM quercetin 81.67 ± 3.18 % NeuN^+^/CldU^+^ cells, n = 3 mice per group, *p* = 0.025; Fig 3G,H), whereas the number of new astrocytes was unchanged (0 μM quercetin 2.33 ± 0.88 vs 50 μM quercetin 0.98 ± 0.008 % S100β^+^/NeuN^+^ cells, n = 3 mice per group, *p* = 0.2). These results indicate that quercetin can elicit pro-neurogenic responses *in vivo*. The apparent decrease in proliferation is in line with the idea that quercetin promotes cell cycle exit.

### Quercetin induces endogenous antioxidants and the Akt pathway

In order to further elucidate the underlying molecular mechanisms whereby quercetin exerts an effect on cell survival, we next performed RNA microarray analysis on cells collected 24 hours after growth factor withdrawal in the presence or absence of 25 μM quercetin. As visualized in the volcano plot shown in Fig. 4A, we detected 3900 transcripts for which the expression levels significantly changed following quercetin treatment. The list of top 10 positively regulated genes (Table 1) were enriched for genes related to oxidative stress (Gsta3, Srxn1, Osgin1), as well as Relaxin1, which has been reported to promote osteoblast differentiation (Moon et al., 2014). Remarkably, we found that Miat, a long non-coding RNA recently identified as a switch gene in lineage determined neuronal progenitor cells during cortical development (Aprea et al., 2013) was among the top-10 negatively regulated transcripts. In order to systematically identify putative target pathways we subjected the transcript lists to enrichment with WebGestalt and pathway analysis from the KEGG and WikiPathway databases. Following hypergeometric testing, the top 10 most significantly enriched pathways were identified (Fig 4B,C). The pathways were then ranked according to the ratio of genes in the sample to the overall number of members in that pathway. In both databases, pathways involved in “cell cycle,” “endogenous antioxidative activity,” and “cell survival” were significantly enriched. In both databases mitogen-activated protein kinase (MAPK) signaling was also enriched. It is known that, depending on their specific chemical structure, flavonoids can either inhibit or activate these pathways (reviewed by (Spencer, 2009)). The flavonoids were reported to exert their effect on cell survival through modulation of the MAPK and phosphoinositide-3 kinase (PI3K-Akt) pathways. The enrichment analysis of the MAPK pathway showed that ERK was not differentially expressed in response to quercetin treatment. In contrast, JNK2 and JNK3 were upregulated, whereas JNK1 was downregulated. The transcripts of JNKs-related proapoptotic genes, namely Bid, Bax and PUMA, were also upregulated in the quercetin-treated group (Table 2). To determine whether the pro-survival and anti-apoptotic signals were simultaneously upregulated, we next analyzed the enrichment of the differentially expressed genes in the PI3K/Akt pathway from the KEGG database and the Nrf2-Keap1 pathway from the WikiPathways database. In the PI3K/Akt pathway we observed the significant upregulation of pro-survival genes such as Akt, CREB, MDM2 and Bcl xL (Bcl2l1). In contrast, apoptosis-inducing genes such as p53 and Bim (Bcl2l11) were downregulated (Table 3).

**Table 1.**
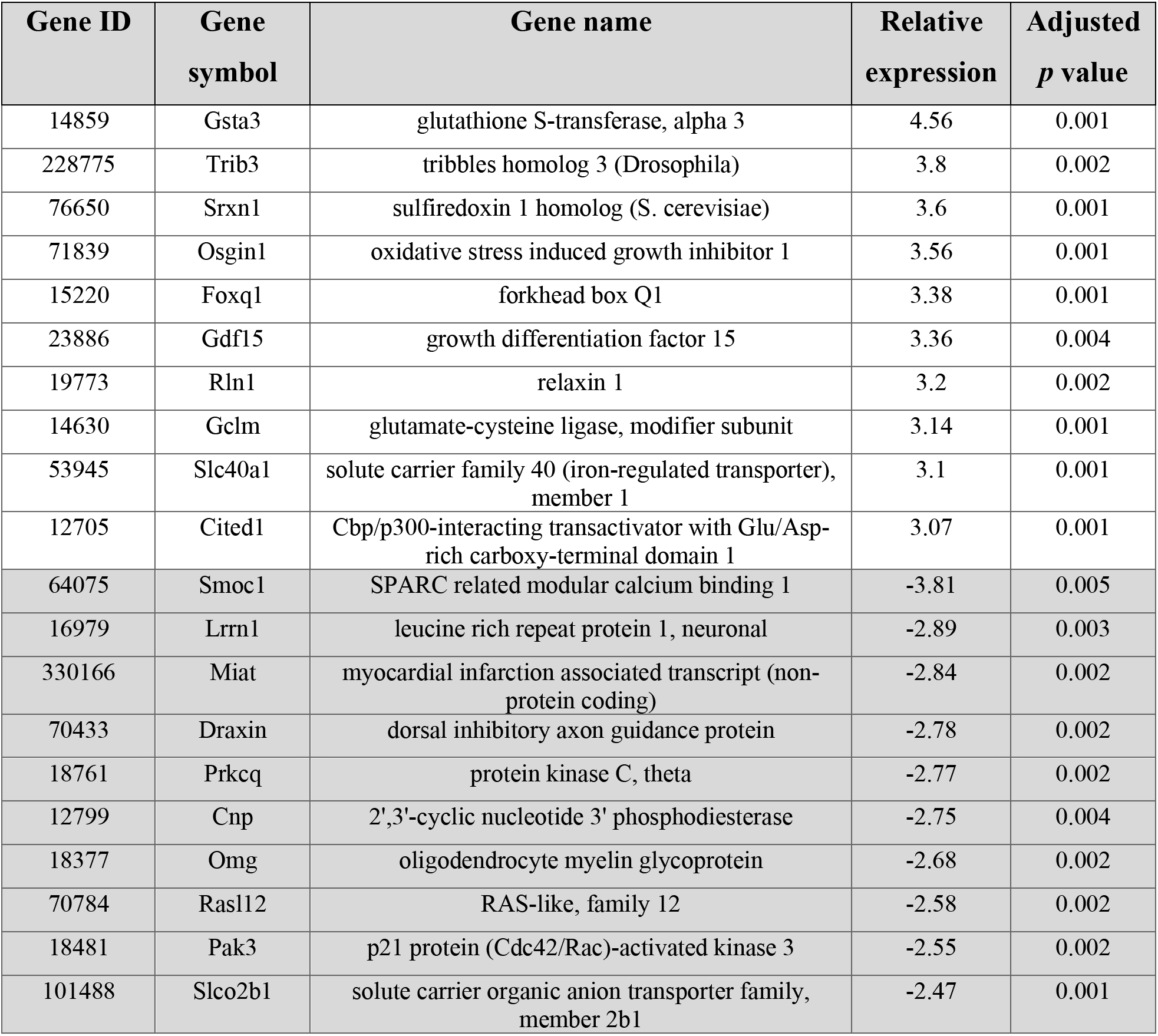
List of the top 10 upregulated and downregulated genes in quercetin-treated adherent neural precursor cells cultured under differentiation conditions.

**Table 2.**
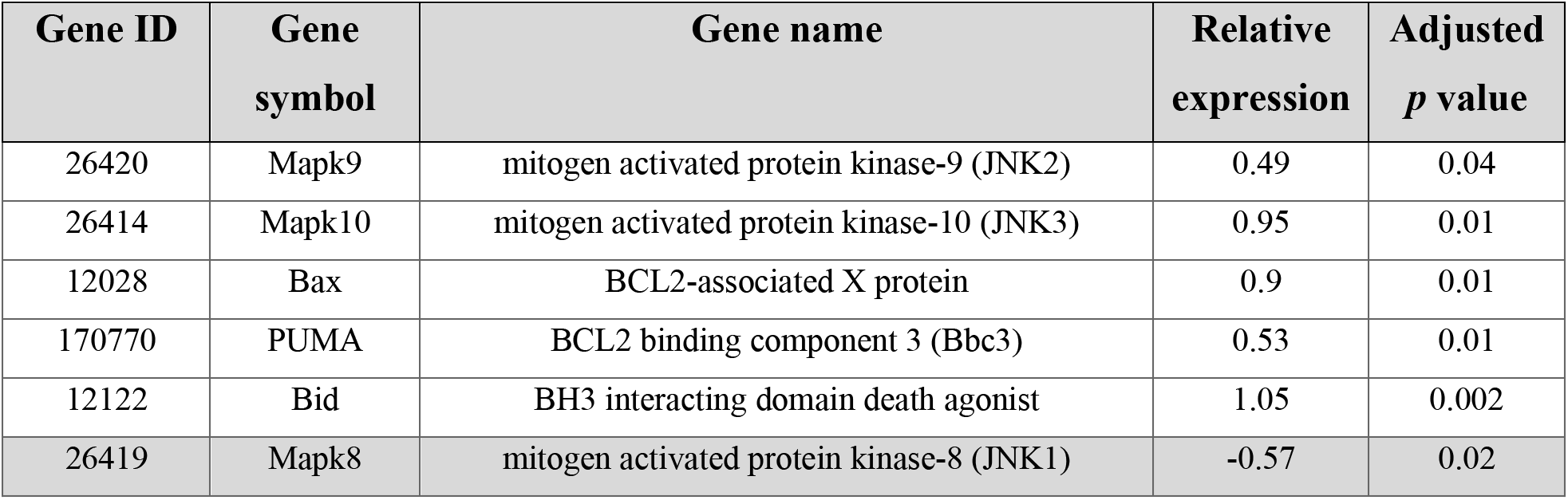
List of the differential expression of JNKs and related pro-apoptotic genes in the MAPK pathway in quercetin-treated adherent neural precursor cells cultured under differentiation conditions.

**Table 3.**
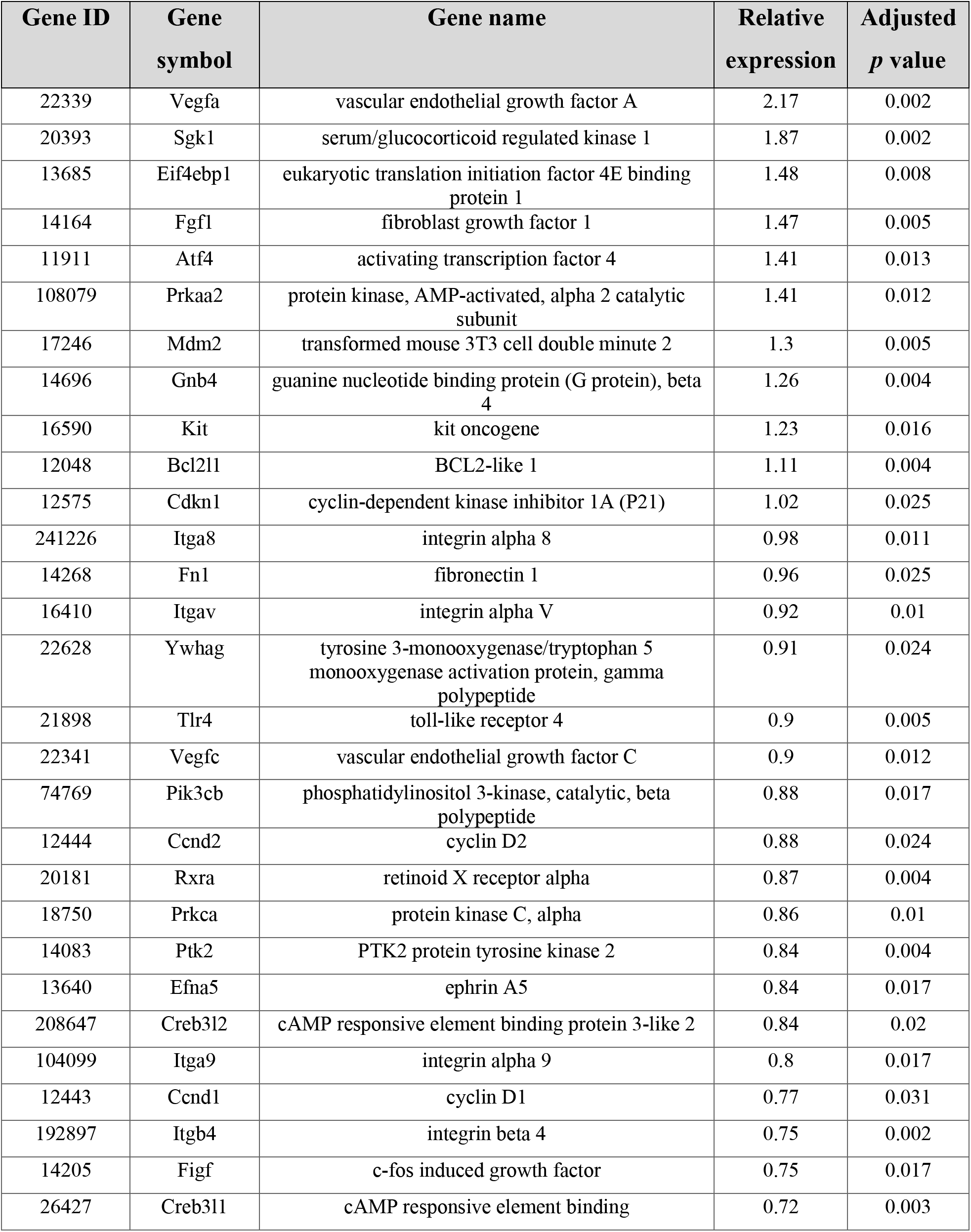
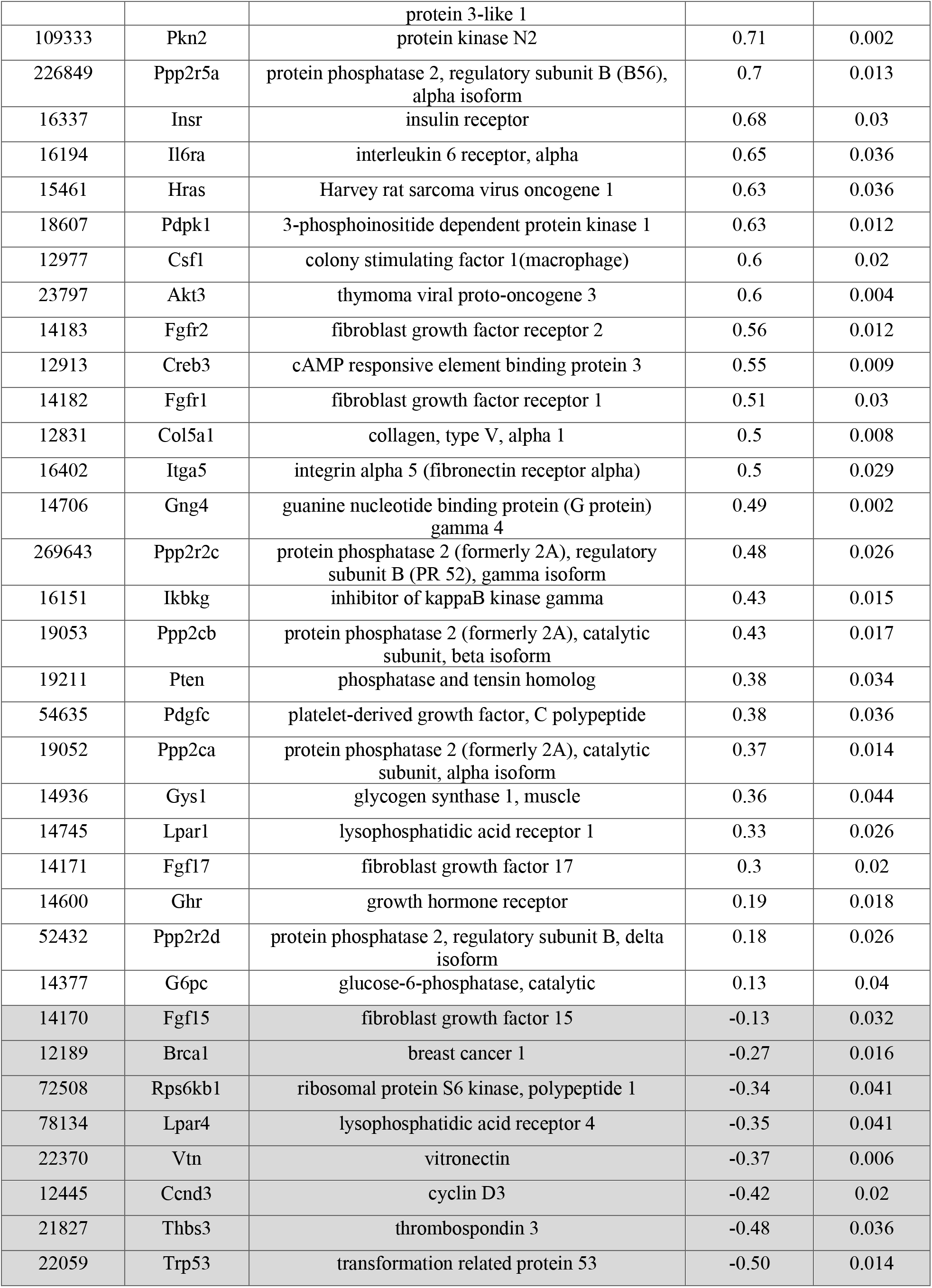
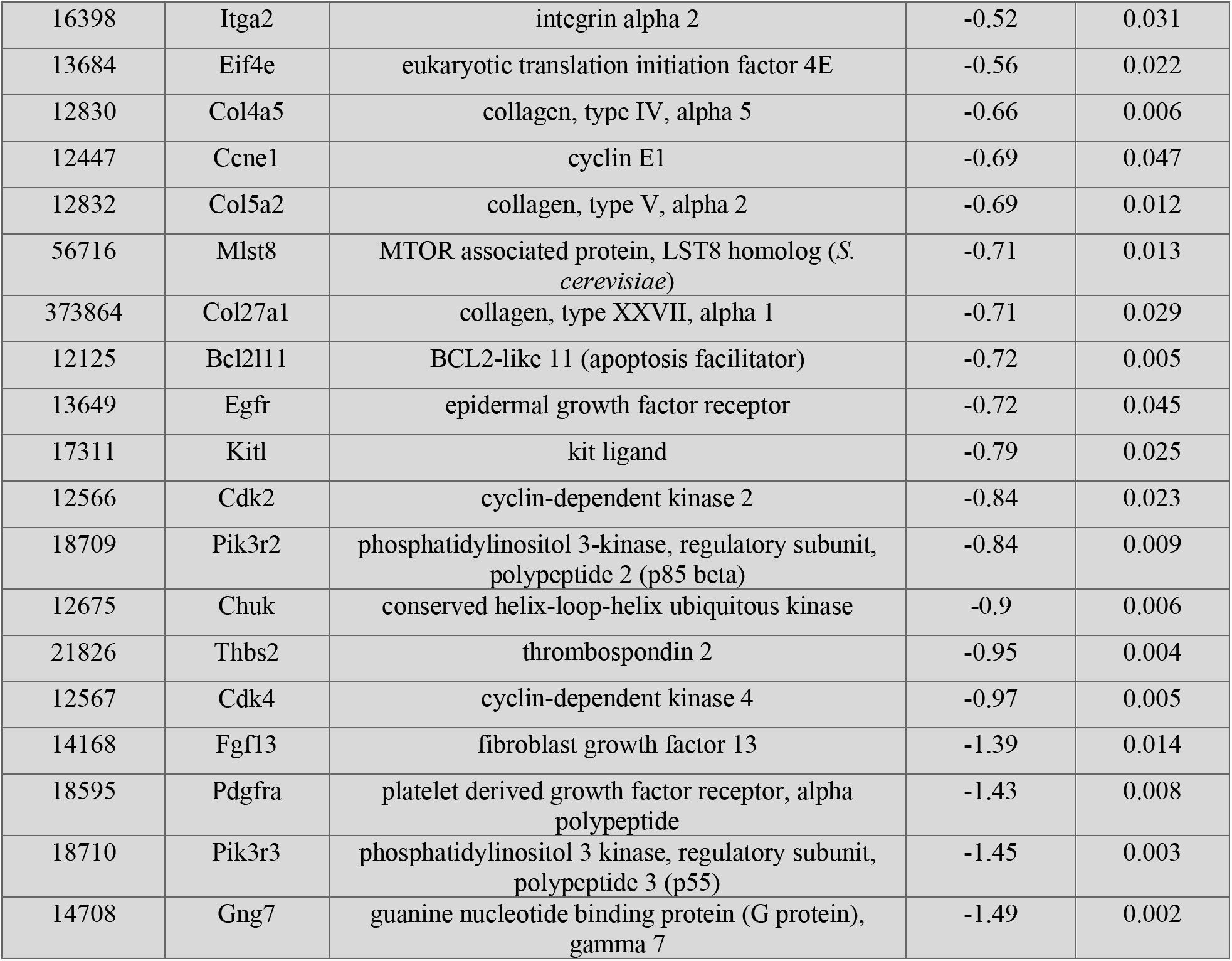
A list of the differentially expressed genes mapped to the PI3 kinase-Akt pathway (KEGG pathways mmu04151) in quercetin-treated adherent neural precursor cells cultured under differentiation conditions.

**Figure 4.**
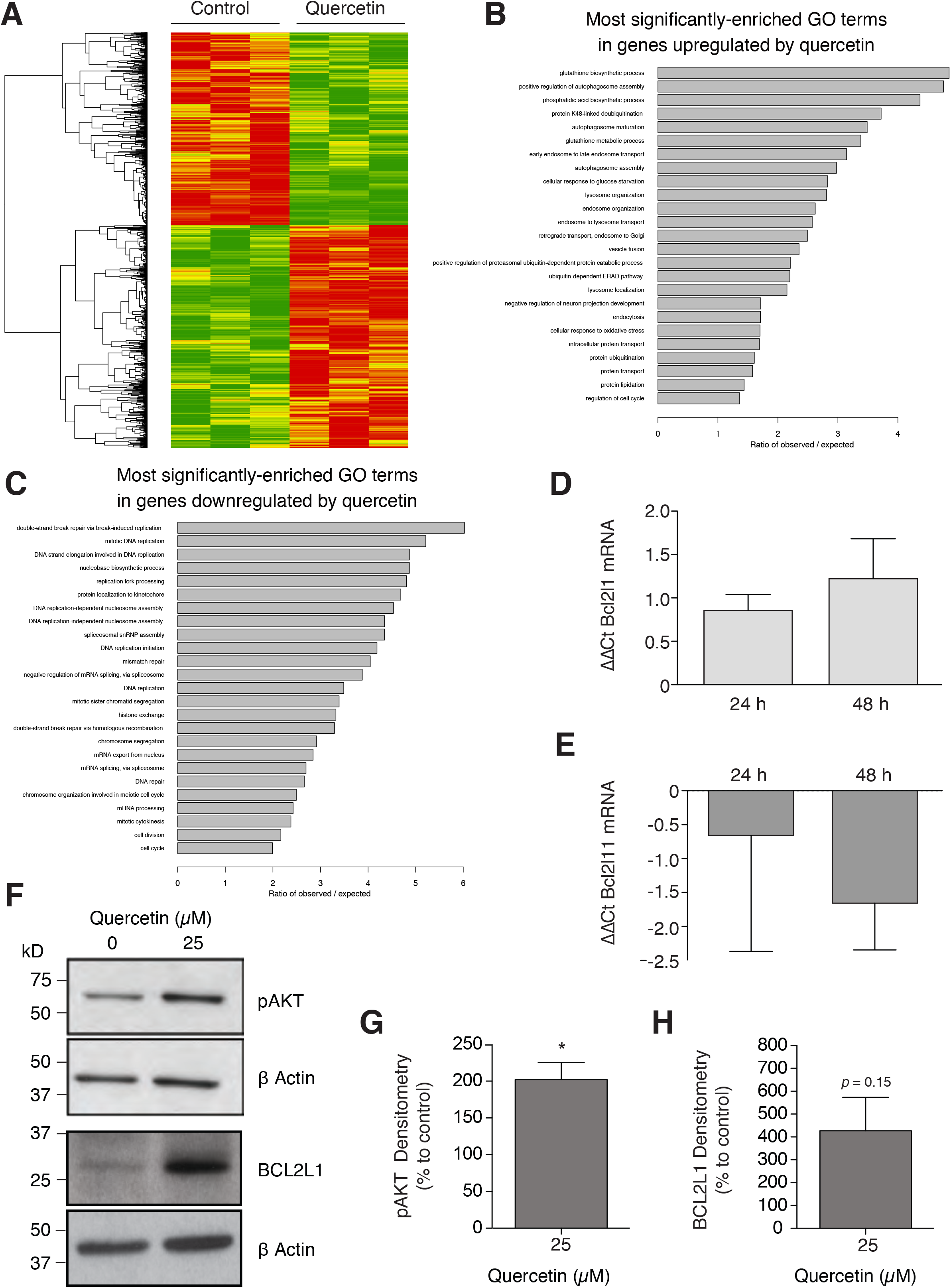
Quercetin induces endogenous antioxidants and the Akt pathway. (A) A heatmap showing changes in neural precursor cell transcript expression in cells treated with 25 μM quercetin, 24 h after growth factor withdrawal. The top 25 most enriched pathways from the KEGG database for significantly upregulated (B) or downregulated (C) transcripts. Relative expression of the Bcl2l1 gene increased (D) and the Bcl2l11 gene decreased (E) in differentiating neural precursor cells after 24 and 48 h quercetin (25 μM) treatment (n = 3 experiments). (F-H) An increase in the expression of pAkt and Bcl2l1 following quercetin treatment was also observed at the protein level. * *p* < 0.05; one sample t-test. Error bars represent SEM.

To confirm the transcriptomic data, we performed qPCR to quantify the expression level of Bcl xL (Bcl2l1) and Bim (Bcl2l11). In line with the microarray data, we found that Bcl2l1 was upregulated over time in the quercetin-treated group (ΔΔCt 1.85 ± 0.2 and 2.55 ± 0.67 after 24 and 48 h of treatment, respectively (Fig 4D). In contrast, the expression of the pro-apoptotic gene Bcl2l11 was downregulated (ΔΔCt −1.55 ± 3.6 and −3.5 ± 1.5 after 24 and 48 h treatment, respectively (Fig 4E). The increased expression of phospho-Akt (pAkt) and Bcl2l1 was also observed at the protein level. After 48 h incubation with 25 μM quercetin, pAKT expression was significantly upregulated (202 ± 23.29 % of control, n = 4, *p* = 0.02; Fig 4F,G). A similar pattern was also observed in BCL2l1 protein expression (427.2 ± 145.6 % of control, n = 3, *p* = 0.15; Fig 4F,H). Another highly enriched pathway that potentially explains the mechanism via which quercetin promotes the survival of differentiating neural precursor cells is the Nrf2-Keap1 pathway. One of the downstream genes in this pathway, Gsta3 (glutathione S-transferase alpha 3), was the most highly expressed gene in differentiating neural precursor cells upon 25 μM quercetin treatment (Table 4). In addition to Nrf2 uprgulation, the Nrf2 antagonist Keap1 (kelch-like ECH-associated protein 1) was also increased in the quercetin-treated group. Therefore, the increase in Keap1 expression upon quercetin treatment may counteract the detrimental effect of excessive Nrf2 accumulation.

**Table 4.** A list of the differentially expressed genes mapped to the Nrf2-Keap1 pathway (Wikipathways WP1245) in quercetin-treated adherent neural precursor cells cultured under differentiation conditions.

| Gene ID | Gene symbol | Gene name | Relative expression | Adjusted *p* value |
| --- | --- | --- | --- | --- |
| 14859 | Gsta3 | glutathione S-transferase, alpha 3 | 4.56 | 0.001 |
| 12608 | Cebpb | CCAAT/enhancer binding protein (C/EBP), beta | 1.96 | 0.004 |
| 15368 | Hmox1 | heme oxygenase (decycling) 1 | 2.24 | 0.003 |
| 17132 | Maf | avian musculoaponeurotic fibrosarcoma (v-maf) AS42 oncogene homolog | 0.36 | 0.033 |
| 18104 | Nqo1 | NAD(P)H dehydrogenase, quinone 1 | 1.91 | 0.002 |
| 18750 | Prkca | protein kinase C, alpha | 0.86 | 0.01 |
| 14630 | Gclm | glutamate-cysteine ligase, modifier subunit | 3.14 | 0.001 |
| 50868 | Keap1 | kelch-like ECH-associated protein 1 | 0.53 | 0.01 |
| 14629 | Gclc | glutamate-cysteine ligase, catalytic subunit | 0.87 | 0.006 |
| 18024 | Nfe2l2 | nuclear factor, erythroid derived 2, like 2 | 0.34 | 0.03 |
| 26419 | Mapk8 | mitogen-activated protein kinase 8 | -0.57 | 0.021 |

### Apple peel and flesh are pro-neurogenic in vitro

Given our findings regarding the flavonoid quercetin as a potential pro-neurogenic compound, we next wanted to determine whether apple peel or flesh display similar pro-neurogenic properties. We first sought to identify an apple variety (cultivar) that is particularly rich in flavonoids. As Saxony, where our institutes are located, has a long tradition of breeding and growing apples, we compared six locally harvested varieties: Elstar, Pinova, Pilot, Rebella, Roter Berlepsch and Jonagold. Using the borohydride-chloranil assay, we found that these varieties differed greatly in their absolute flavonoid contents, but all contained flavonoids in their peel, and to a much lesser extent in their flesh (Table 5). The enrichment in the peel was greatest (with a factor of almost 9-fold) in the Pinova cultivar (Malus domestica ‘Pinova’). Based on these findings we chose Pinova (also marketed as Pinata, Sonata, or Corail) for subsequent experiments. Peel and flesh were extracted separately the solvent removed and the extract dissolved in DMSO. Using the neurosphere assay, we found that Pinova peel (140.1 ± 14.73 % of control neurospheres, n = 3 experiments, *p* < 0.05; Fig 5A) and flesh extract (156.8 ± 8.67 % of control neurospheres, n = 3 experiments, *p* < 0.001; Fig 5A) had a similar pro-neurogenic capacity. In absolute terms, the effect was mild, but there was no difference between flesh and peel as one might have predicted from the different abundance of flavonoids in the two compartments.

**Table 5.** The total flavonoid content in different apple varieties.

| **Cultivar** | **Total flavonoid content**<br>**(equivalent to mg quercetin / 100 g fresh weight)** |  |
| --- | --- | --- |
|  | **Peel** | **Flesh** |
| Pinova | 379 ± 93.95 | 42.45 ± 17.63 |
| Elstar | 428.9 ± 103.7 | 96.74 ± 32.61 |
| Pilot | 344.3 ± 74.19 | 67.36 ± 25.43 |
| Rebella | 186.6 ± 44.4 | 48.82 ± 15.10 |
| Roter Berlepsch | 148.1 ± 60.55 | 18.74 ± 7.86 |
| Jonagold | 87.85 ± 28.45 | 36.98 ± 15.53 |

**Figure 5.**
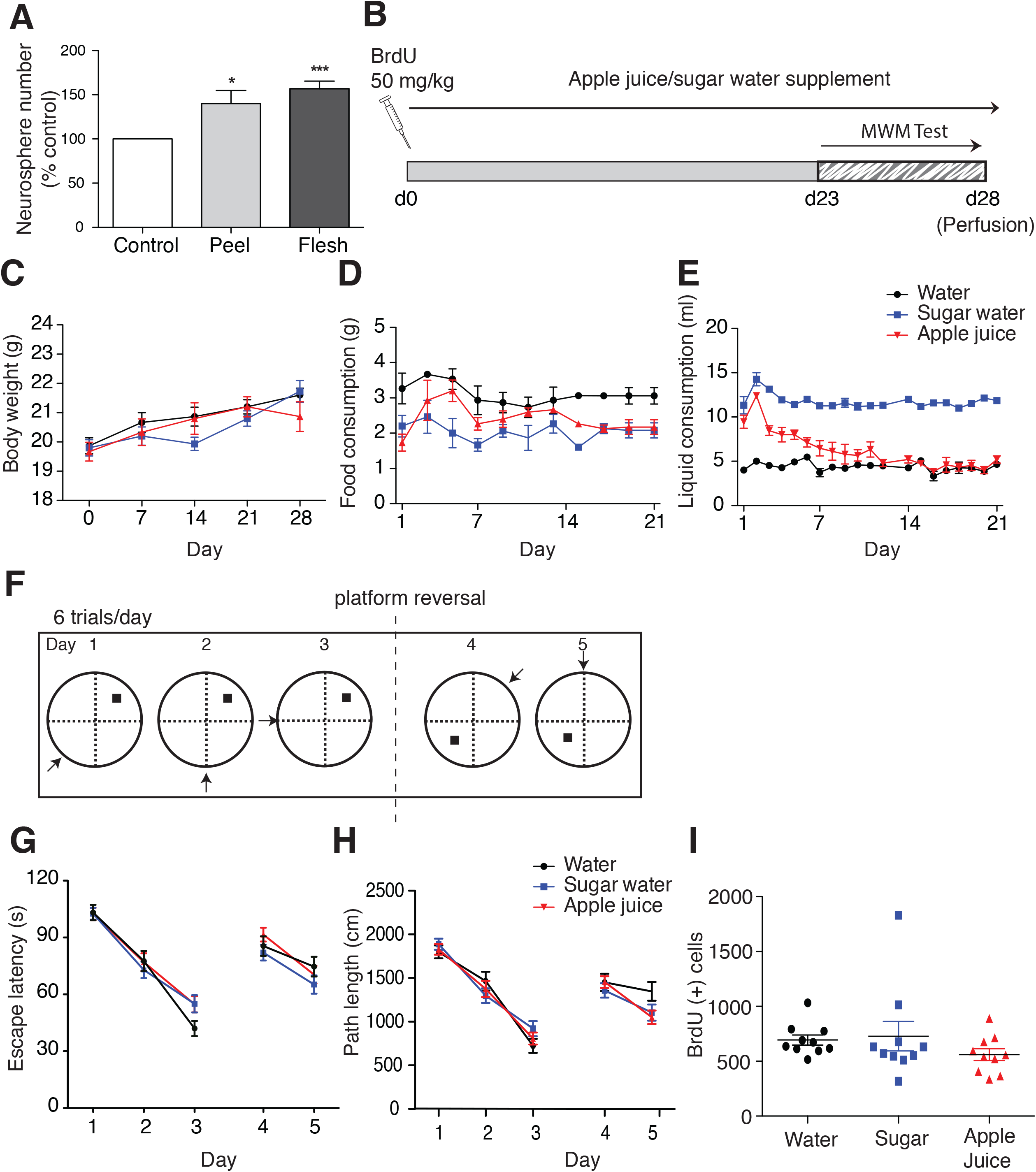
Apple peel and flesh are pro-neurogenic *in vitro*, but apple juice does not affect *in vivo* neurogenesis or cognitive function. (A) Pinova apple peel and flesh extracts significantly increased the number of primary hippocampal neurospheres. * *p* < 0.05, *** *p* <0.001; one-way ANOVA with Dunnett’s post-hoc test. (B) Experimental scheme for apple juice supplementation. (C) No significant difference in body weight was observed between treatment groups. (D) The water-supplemented group consumed significantly more food than the apple juice- and sugar water-supplemented mice. (E) The sugar water-supplemented mice consumed significantly more liquid than the other groups. (F) Experimental scheme for the performance of the Morris water maze task. No effect of apple juice or sugar water was observed measured by the escape latency (G) or the path length (H). No difference in the number of BrdU^+^ cells was observed between treatment groups. One-way ANOVA. Error bars represent SEM.

### Apple juice supplementation does not affect adult neurogenesis in vivo

Given that apple juice is one of the most widely consumed fruit beverages, we next examined whether consumption of whole apple juice concentrate affected adult neurogenesis *in vivo,* with concomitant effects on learning and memory (Fig 5B). Although the *in vitro* data suggested only small responses, we did not want to miss an interesting effect. We also wanted to exclude the possibility that a specific effect might be masked by systemic factors not present in the reductionist setting of the culture dish.

To exclude the effect of the increased caloric intake of fruit sugar as a potential confounding factor, a group that received equicaloric sugar water was included in addition to the control group that received normal drinking water. However, all mice showed a similar change in body weight over the experimental period (Fig 5C). Food consumption was greater in mice that drank water compared to mice which received apple juice or equicaloric sugar water (Fig 5D). Liquid consumption differed greatly between the groups, with the highest consumption in the sugar water group (Fig 5E). In order to detect whether the apple juice supplement had an effect on hippocampus-dependent learning, a reversal version of the Morris water maze task, designed to detect the specific contribution of adult-generated neurons to the overall performance in spatial navigation and cognitive flexibility, was performed (Garthe, Behr, & Kempermann, 2009; Garthe, Huang, Kaczmarek, Filipkowski, & Kempermann, 2014; Garthe & Kempermann, 2013); Fig 5F). We did not detect any difference between the groups with respect to escape latency (Fig 5G), length of swim path (Fig 5H), probe trial performance or reversal learning (Fig 5G,H). Following the behavioral testing, the mice were perfused and the number of surviving BrdU^+^ cells was quantified. No effect of either the sugar water or apple juice supplementation on adult hippocampal neurogenesis was observed (n = 10 animals per group, *p* = 0.375; Fig 5I).

### Pro-neurogenic activity in apple flesh

We next returned to our earlier observation that, despite the lower level of flavonoids in apple flesh, there was still as much pro-neurogenic activity as in peel extract. To identify additional pro-neurogenic compounds in apple extracts we used a bioassay-guided fractionated approach. Flesh extract was separated into fractions using liquid- and solid phase separation methods (Fig 6A). The Pinova flesh extract was dissolved in an equal mixture of water and ethyl acetate (EtOAc) to separate the polar from the non-polar compounds. The water fraction 1 was further sub-fractionated by addition of an equal volume of butanol (BuOH) to obtain water fraction 2 and a BuOH fraction. The “water fraction” referred to in the remainder of the text is water fraction 2. Among the three fractions (water, EtOAc and BuOH), we found a dose-dependent increase in the number of neurospheres generated from primary dentate gyrus cell cultures in the presence of the water fraction, more than doubling the number of neurospheres (1 mg/ml 227.8 ± 24.86 %, 3 mg/ml 235.8 ± 17.12 % of control neurospheres, n = 5 experiments, *p* < 0.001; Fig 6B). The EtOAc fraction decreased neurosphere counts and there was a small increase in the BuOH fraction, which could not be further explored due to the low yield. As the water fraction also contains fruit sugar that might affect cell growth, we tested a solution with a sugar composition (sucrose, glucose and fructose) equivalent to the water fraction. There was no effect of sugar on neurosphere numbers (Fig 6C).

**Figure 6.**
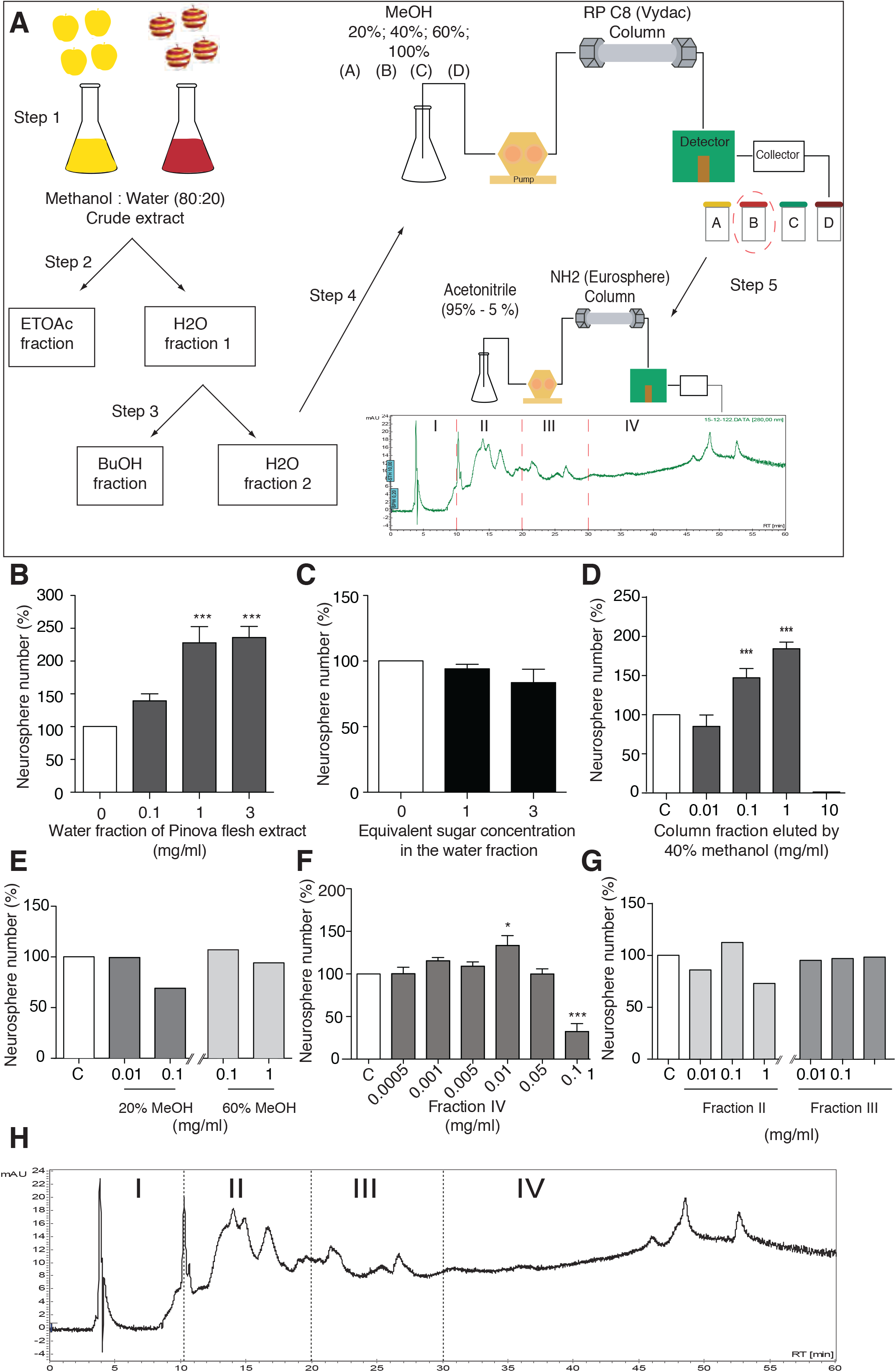
Apple flesh extracts display pro-neurogenic activity. (A) Experimental scheme for the extraction and fractionation of apple extract. Apple flesh was extracted with 80% methanol in water (step 1). Nonpolar compounds were enriched in EtOAc (step 2). Less polar compounds were enriched in BuOH (step 3). The polar compounds in the water fraction were further fractionated by column fractionation using a C8 column in reverse phase HPLC system (step 4). The active fraction from step 4 (fraction B, eluted in 40% MeOH) was subsequently fractionated using an NH2 column. (B) The water fraction of Pinova flesh extract significantly increased the number of hippocampal neurospheres, whereas the equivalent concentration of sugar had no effect (C). Following further separation of the water fraction, the 40 % methanol-eluted fraction significantly increased the number of hippocampal neurospheres (D), whereas the 20 % and 60 % methanol fractions had no effect (E). Following further fractionation of the 40 % methanol fraction, the pro-neurogenic activity was restricted to fraction IV (F-H). * *p* < 0.05, *** *p* < 0.001; one-way ANOVA with Dunnett’s post-hoc test. Error bars represent SEM.

The water fraction was next further fractionated by HPLC in solvents of different polarity (20% to 100% methanol, MeOH, in water). The 40% MeOH eluent increased neurosphere number in a dose-dependent manner (85.09 ± 7.3 %; 147.2 ± 5.88 %; and 184.4 ± 4.18 % of control neurospheres for 0.01, 0.1, and 1 mg/ml, respectively, n = 4 experiments, *p* < 0.001; Fig 6D), but decreased neurosphere counts at the high concentration of 10 mg/ml. The fractions eluted at 20% and 60% MeOH did not increase neurosphere numbers (Fig 6E) and no fraction could be obtained at 100% MeOH. The 40% MeOH fraction was further fractionated using a NH_2_ column, in which hydrophilic compounds are eluted first. Acetonitrile was used as eluent, again graded from 80% down to 5% acetonitrile. Four fractions were distinguished (Fig 6F-H). The neurosphere assay revealed that fraction IV dose-dependently increased the neurosphere count, with a maximum effect at a concentration of 0.01 mg/ml (133.5 ± 11.46 % of control neurospheres, n = 6 experiments, *p* < 0.05; Fig 6F). At higher concentrations the number of neurospheres was again decreased. No effect of fraction II or III was observed in terms of neurosphere number (Fig 6G).

Fraction IV could not be further sub-fractionated because of the low amount of material. We instead used ultra-performance liquid chromatography coupled with photodiode-array detector and electrospray quadrupole time-of-flight mass spectroscopy (UPLC/PDA/ESI-qTOF-MS) to identify the compounds present in this fraction. The compounds eluted in the first 1.5 min could not be separated, but subsequent peaks (numbers 1-6; Fig 7A) were subjected to TOF-MS and collision-induced dissociation (CID). Interestingly, two of the peaks were identified as the glycosides of dihydroxybenzoate (Table 6). Dihydroxybenzoate is the salt of dihydroxybezoic acid (DHBA), which belongs to the non-flavonoid phenolic compounds present in fruits. We therefore obtained commercially available isomers of DHBA and examined their effect on neural precursor cell activity. A significant increase in neurosphere numbers was found with 2,3-DHBA at a concentration of 1 μM (115.9 ± 4.69 % of control neurospheres, n = 5 experiments, *p* < 0.05; Fig 7B) and 3,5-DHBA at concentrations of 1 μM and 10 μM (10 μM 161.12 ± 14.97 % of control neurospheres, n = 5 experiments, *p* < 0.001; Fig 7C). Treatment with 1 μM 3,4-DHBA produced a trend towards an increase in the number of neurospheres generated, but this failed to reach statistical significance (*p* = 0.19; Fig 7C). We repeated the neurosphere experiment with 10 μM 3,5-DHBA and confirmed the above results, observing a significant increase in the number of neurospheres generated from the 3,5-DHBA-treated cells (control: 26.75 ± 3.5 vs 3,5-DHBA: 38.75 ± 2.72 neurospheres, n = 4 experiments, *p* = 0.03). Treatment with 10 μM 3,5-DHBA also resulted in a significant increase in the size of the neurospheres (control 53.52 ± 2.7 μm vs 3,5-DHBA: 71.53 ± 3.9 μm diameter, n = 4 experiments, *p* = 0.0007; Fig 7E). We also found that treatment of adherent monolayer cultures with 3,5-DHBA at concentrations of 100 μM and 250 μM resulted in a higher cell density after 4 days in culture (0 μM 3.16 ± 0.06 vs 100 μM 3.78 ± 0.16 vs 250 μM 3.88 ± 0.17, n = 8 experiments, *p* = 0.0001; Fig 8A). To determine whether this was due to 3,5-DHBA acting on precursor cell survival when cultured in proliferation conditions, adherent cells were cultured in growth factors for 4 days and varying concentrations of 3,5-DHBA were added on either day 0 (4 day), day 2 (2 day), or day 3 (1 day), after which the Resazurin cell viability assay was performed on day 4 (Fig 8B). We found an increase in the percentage of cell survival at the higher 3,5-DHBA concentrations, with the strongest effect size observed when the 3,5-DHBA was present in the medium for the entire 4 days of the experiment (Fig 8C-E). We also examined whether 3,5-DHBA had a similar cell survival effect when the adherent precursor cells were cultured under differentiation conditions (Fig 8F). Again we observed a neuroprotective effect, with a significant increase in cell survival found in the cells treated with 3,5-DHBA (≥ 250 μM) throughout the entire differentiation process (250 μM 3,5-DHBA 113.5 ± 2.5 % of control, n = 6 experiments, *p* = 0.0005; Fig 8G). After 6 days of differentiation these cells were fixed and stained for markers of mature neurons and astrocytes. We found a significantly higher percentage of βIII-tubulin^+^ neurons in the cultures differentiated in the presence of high concentrations of 3,5-DHBA, further confirming its pro-neurogenic effect (0 μM 3,5-DHBA 25.42 ± 3.1 vs 1000 μM 3,5-DHBA: 90.54 ± 2.3 % βIII-tubulin^+^ cells, n = 5 experiments, *p* <0.0001; Fig 8H, I).

**Table 6.** Peak annotation of UPLC / PDA / ESI-QTOFMS analysis of fraction IV based on accurate mass measurement and high-resolution CID mass spectra.

| Peak number | *m/z*, ESI(+) | *m/z*, ESI(-) | Elemental composition | Comments |
| --- | --- | --- | --- | --- |
| 1 | 347.0942, [M<sup>+</sup>Na]<sup>+</sup> | 323.0979, [M<sup>-</sup>H]<sup>-</sup> | C<sub>12</sub>H<sub>20</sub>O<sub>10</sub> | hexoside |
| 2 | 339.0668, [M<sup>+</sup>Na]<sup>+</sup> | 315.0713, [M<sup>-</sup>H]<sup>-</sup> | C<sub>13</sub>H<sub>16</sub>O<sub>9</sub> | dihydroxybenzoate O-hexoside |
| 3 | 361.1099, [M<sup>+</sup>Na]<sup>+</sup> | 337.1133, [M<sup>-</sup>H]<sup>-</sup> | C<sub>13</sub>H<sub>22</sub>O<sub>10</sub> | hexoside, homolog of 1 |
| 4 | 471.1082, [M<sup>+</sup>Na]<sup>+</sup> | 447.1132, [M<sup>-</sup>H]<sup>-</sup> | C<sub>18</sub>H<sub>24</sub>O<sub>13</sub> | dihydroxybenzoate *O*-(hexoside-pentoside) |
| 5 | 361.1091, [M<sup>+</sup>Na]<sup>+</sup> | 337.1140, [M<sup>-</sup>H]<sup>-</sup> | C<sub>13</sub>H<sub>22</sub>O<sub>10</sub> | hexoside, Isomer of 3 |
| 6 | 660.1686, [M<sup>+</sup>H]<sup>+</sup> | 658.1546, [M<sup>-</sup>H]<sup>-</sup> | ? | ? |

**Figure 7.**
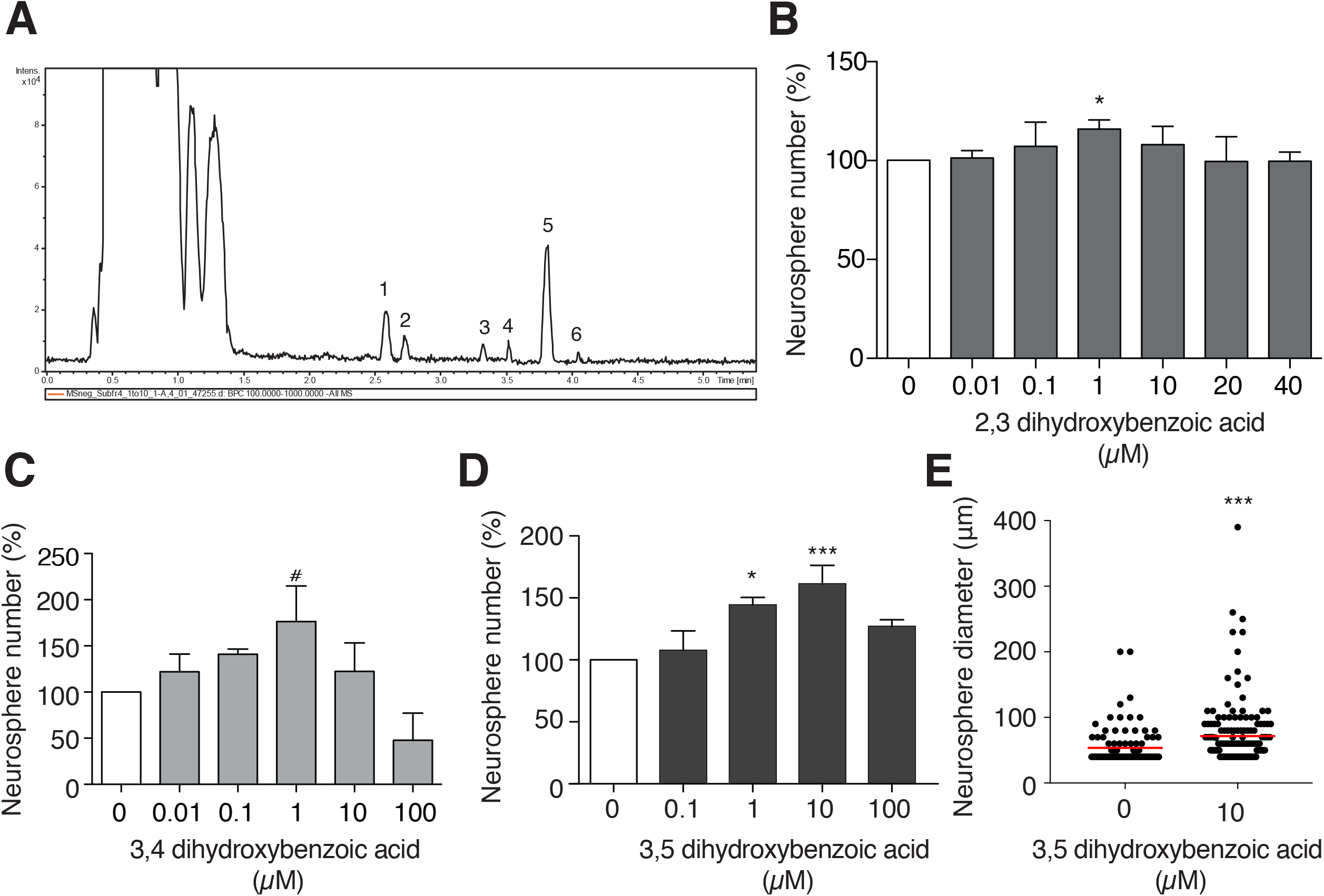
Identification of hydroxybenzoic acid is a pro-neurogenic compound found in apple flesh. (A) Base peak chromatogram m/z 100-1000, gradient elution, ESI(-) of fraction IV. Six peaks were annotated, two of which (peaks 2 and 4) were glycosides of dihydrobenzoate. Neurosphere assays revealed that the dihydroxybenzoic acid isomers (B) 2,3-DHBA, (C) 3,4-DHBA and (D) 3,5-DHBA all increased precursor proliferation, with the greatest effect observed in response to 3,5-dihydroxybenzoic acid. (E) In addition to increasing neurosphere number, 3,5-DHBA also significantly increased the diameter of the resulting neurospheres. * *p* < 0.05, *** *p* < 0.001; one-way ANOVA with Dunnett’s post-hoc test (B, C and D) and Student’s t-test (E). Error bars represent SEM.

**Figure 8.**
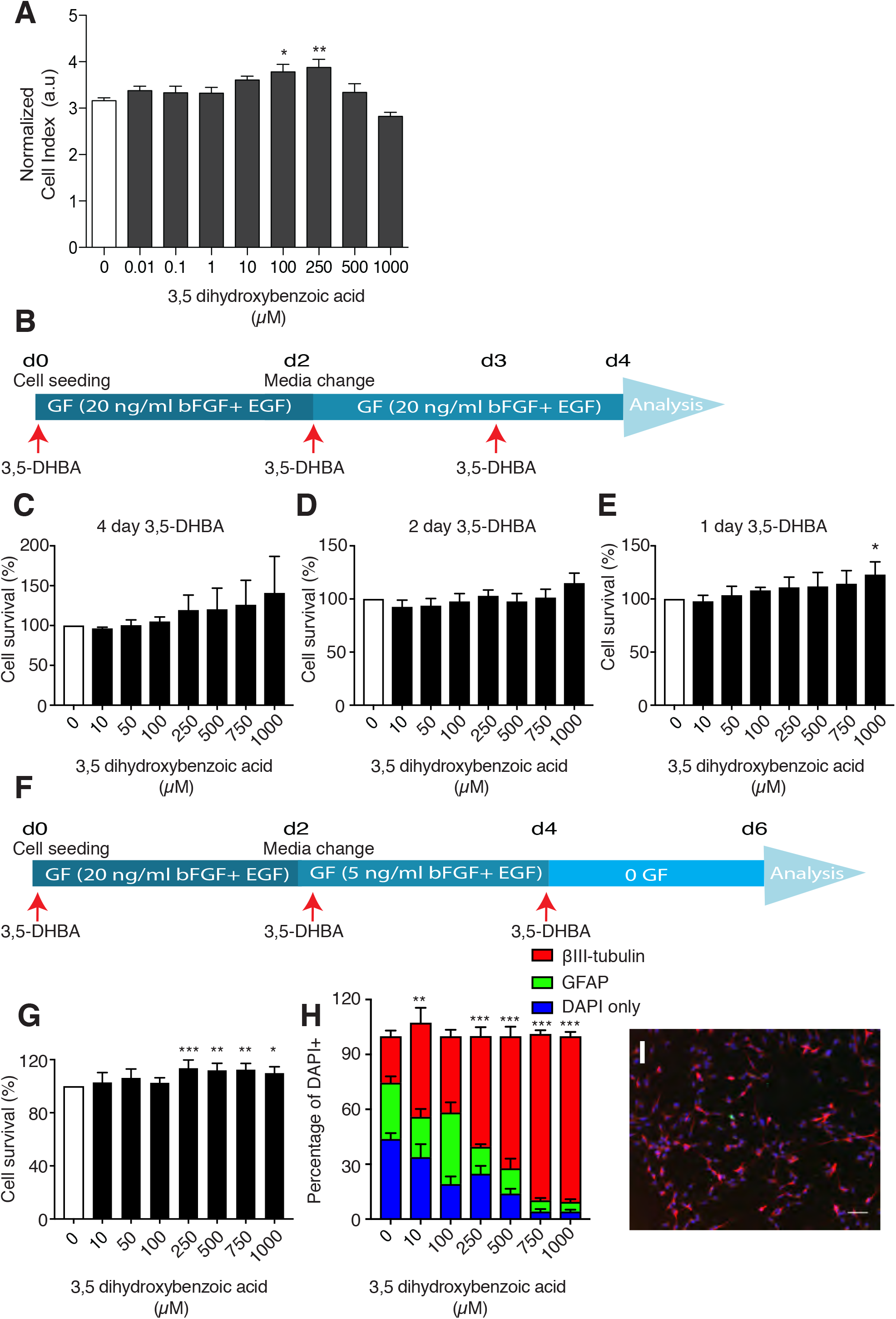
DHBA promotes *in vitro* cell survival and neuronal differentiation. (A) Using the xCELLigence impedance-based assay, an increased number of cells were detected in adherent monolayer cultures treated with 100 and 250 μM 3,5-DHBA. (B) Adherent cells were cultured in 20 ng/ml EGF and bFGF for 4 days and varying concentrations of 3,5-DHBA were added on either day 0 (d0), day 2 (d2) or day 3 (d3). The Resazurin cell viability assay was then performed on day 4. (C-E) An increase in the percentage of cell survival at the higher 3,5-DHBA concentrations was observed. (F) Adherent precursor cells were cultured under differentiation conditions with 3,5-DHBA present in the culture medium throughout the experiment. (G) A significant increase in cell survival was observed in the differentiating cells treated with ≥ 250 μM 3,5-DHBA. (H) 3,5-DHBA dose-dependently increased the number of surviving differentiated neurons, with the highest effect observed in the cultures treated with 1000 μM 3,5-DHBA (I). * *p* < 0.05, ** *p* < 0.01, *** *p* < 0.001; one-way ANOVA with Dunnett’s post-hoc test. Error bars represent SEM. Scale bar in I is 50 μm.

### HCAR1, the putative receptor for hydroxybenzoic acid, is expressed on neural precursor cells and in the neurogenic niche in vivo

The hydroxycarboxylic acid receptor 1 (HCAR1) had been listed as orphan receptor until it was identified as the receptor for lactate and 3,5-DHBA. In adipocytes, activation of the receptor inhibits lipolysis (Liu et al., 2012). HCAR1 is also expressed in the brain, including the hippocampus (Castillo et al., 2015). Using immunohistochemistry we confirmed that HCAR1 is expressed by the proliferating Nestin^+^/Sox2^+^ hippocampal precursor cells (Fig 9A). Immunofluorescence staining of brain sections revealed that, in addition to being expressed by the granular cells of the dentate gyrus, as has been previously reported by Castillo et al., 2015, HCAR1 is also expressed by type-1 and type-2 precursor cells, as evidenced by the co-expression of Nestin and Sox2 (Fig 9B).

**Figure 9.**
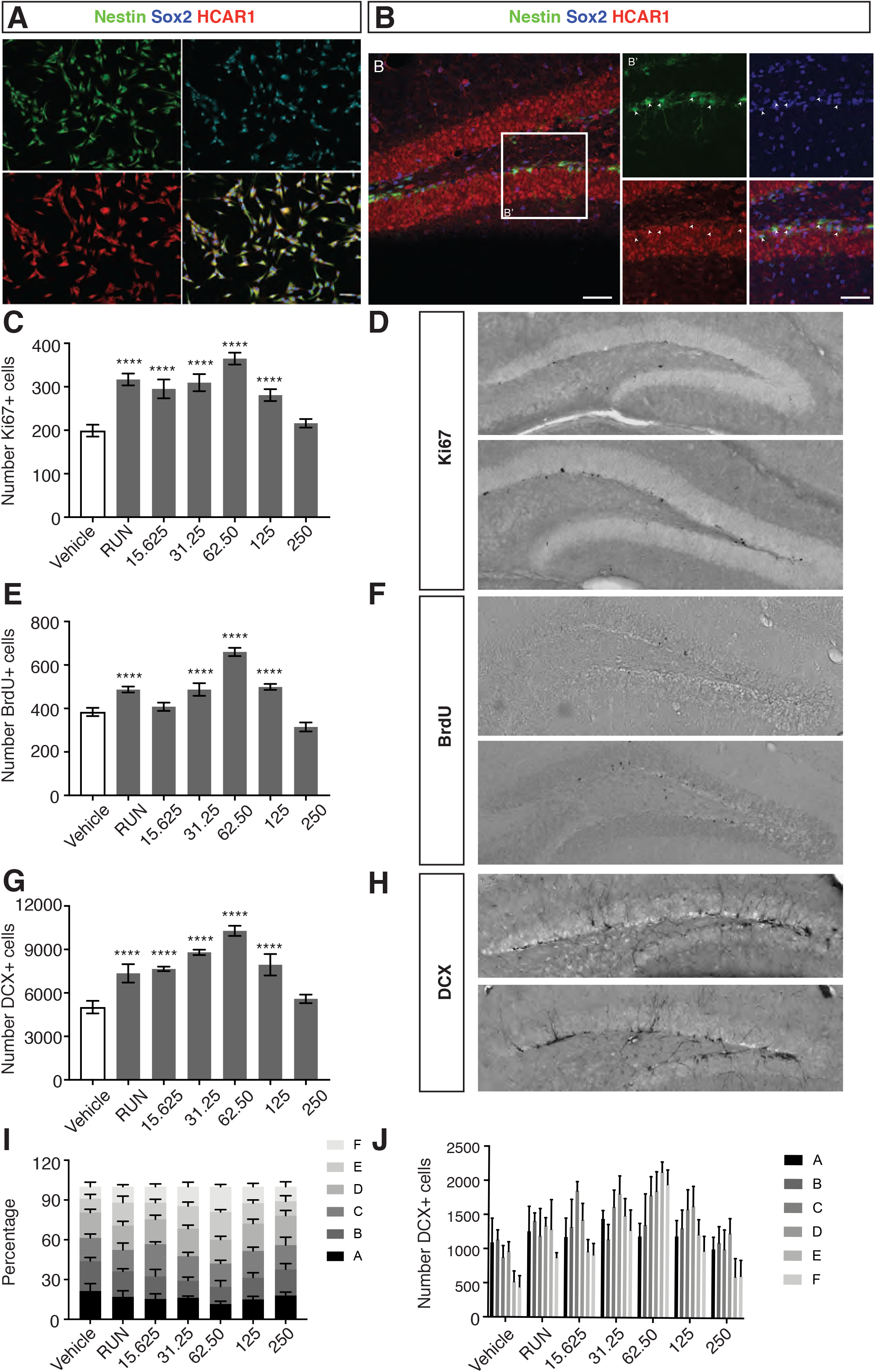
3,5-DHBA increases neural precursor cell proliferation and neurogenesis *in vivo*. (A) Immunofluorescence staining of proliferating adult hippocampal precursor cells in monolayer culture. HCAR1 (red), co-localizes with the neural precursor cell markers nestin (green) and Sox2 (cyan). Confocal images of the hippocampal dentate gyrus in brain sections from Nestin-GFP mice at 20x magnification (B) and 40x magnification (B’). HCAR1 (red), nestin (green) and Sox2 (dark blue). Arrowheads show the cells with HCAR1, Nestin and Sox2 colocalization. *In vivo* administration of 3,5-DHBA resulted in a significant increase in the number of proliferating (Ki67^+^) cells (C), BrdU^+^ cells (E,F) and DCX^+^ immature neurons (G,H) in the dentate gyrus. 3,5-DHBA administration altered the dendritic morphology of the newly born neurons, increasing both the percentage (I) and absolute number (J) of DCX^+^ cells with a more mature phenotype. **** *p* < 0.0001; one-way ANOVA with Dunnett’s post-hoc test. Error bars represent SEM. Scale bars are 50 μm.

### 3,5-DHBA increases neural precursor cell proliferation and neurogenesis in vivo

Given the positive effects of 3,5-DHBA on neural precursor cell proliferation and survival *in vitro,* and the presence of the hydroxybenzoic acid receptor, HCAR1 on neural precursor cells *in vivo*, we finally examined whether administration of 3,5-DHBA affects *in vivo* neural precursor cell proliferation and adult neurogenesis. To do this, 3,5-DHBA (15.625 mg/kg, 31.25 mg/kg, 62.5 mg/kg, 125 mg/kg and 250 mg/kg) was administered intraperitoneally to adult mice once daily for 14 days prior to perfusion. A group that received no DHBA, but had free access to a running wheel was included as a positive control. All mice received two intraperitoneal injections of BrdU (50 mg/kg) spaced 1 h apart prior to the first dose of 3,5-DHBA to label the proliferating cells. Treatment with 3,5-DHBA resulted in a significant increase in the number of proliferating neural precursor cells located in the subgranular zone of the dentate gyrus in all but the group receiving the highest dose of 3.5-DHBA, with the greatest effect being observed at a dose of 62.5 mg/kg (vehicle: 199.2 ± 6.1 vs 62.5 mg/kg 3,5-DHBA: 316.8 ± 6.1 Ki67^+^ cells, n = 5 mice per group, *p* < 0.0001; Fig 9 C,D). A similar result was observed following quantification of the number of BrdU-labeled cells (vehicle: 384 ± 8.5 vs 62.5 mg/kg 3,5-DHBA: 660 ± 8.5 BrdU^+^ cells, n = 5 mice per group, *p* < 0.0001; Fig 9 E,F). Finally, the number of immature neurons (DCX^+^) was also counted to determine whether, in addition to proliferation, 3,5-DHBA treatment also influences adult neurogenesis. Similar to the results obtained for the *in vitro* experiments, we observed a significant increase in the number of newborn neurons in all but the group receiving the highest dose of 3,5-DHBA, with the greatest increase observed in the group receiving 62.5 mg/kg (vehicle: 5023 ± 194.8 vs 62.5 mg/kg 3,5-DHBA: 10286 ± 156.6 DCX^+^ cells, n = 5 mice per group, *p* < 0.0001; Fig 9 G,H). To examine whether 3,5-DHBA treatment also affected the maturation rate of the newborn neurons, the DCX^+^ cells were morphologically categorized according to the presence and shape of their apical dendrites, as described in our previously published protocol (Plumpe et al., 2006). Briefly, category A and B cells had no or very short processes, C and D cells had increasingly longer processes that by stage D had reached the molecular layer, category E cells had one strong dendrite with sparse branching, whereas in category F cells the dendritic tree was more elaborate, with multiple branches in the molecular cell layer. We found that the distribution of the DCX^+^ cells into these six phenotypical categories was significantly influenced by 3,5-DHBA treatment. Most notably, the mice treated with 62.5 mg/kg 3,5-DHBA had both a significantly higher percentage and absolute number of DCX^+^ cells belonging to categories E and F than the vehicle-treated control mice, indicating that 3,5-DHBA treatment increases the maturation speed of the newly born neurons (Fig 9 I,J). Interestingly, and consistent with our previous report (Plumpe et al., 2006), this change in dendritic morphology was not observed in the positive control group of mice that exercised.

## Discussion

In this study we used a two-pronged approach to reveal that apples contain pro-neurogenic compounds in both their peel and flesh. We chose a popular local cultivar, Pinova, a cross of Golden Delicious and Clivia, which is very rich in flavonoids with a large peel-to-flesh ratio. We confirm that quercetin, the most prominent flavanol in apple peel indeed stimulates adult hippocampal neurogenesis, both *in vitro* and *in vivo*. Our data confirms a pattern that had already been suggested by the literature. Whereas flavonoids are anti-proliferative at high concentrations (Abosalem, Gibriel, Al-Awady, & Mandour, 2020; Ammendola et al., 2020) and thus have certain anti-carcinogenic properties, at lower concentrations their effect on precursor cells is more complex and appears to be activating rather than inhibitory. Indeed, we found that 25 μM quercetin increased cell cycle exit and promoted survival and differentiation of adult hippocampal precursor cells, both *ex vivo* and *in vivo*. This is similar to the effect that we and others have previously reported for compounds such as resveratrol and EGCG on neural precursor cells (Ortiz-Lopez et al., 2016; Torres-Perez et al., 2015; Yoo et al., 2010). Natural compounds like these, which are abundant in widely consumed fruits and vegetables, might thus exert a small beneficial effect on sustaining neurogenesis and brain plasticity. While consuming the fruits is by themselves not sufficient to promote neurogenesis *in vivo*, the pro-neurogenic effects might be auxiliary and part of environmental influences on brain health.

Based on transcriptomic data we propose that the PI3K-Akt and Nrf2-Keap1 pathways are involved in such effects. This is in agreement with previous studies which indicated that flavonoids have in fact numerous modes of action. During stress conditions, there is an activation and accumulation of Nrf2 in the nucleus (Nguyen, Nioi, & Pickett, 2009). The heterodimer of Nrf2 with small Maf (musculoaponeurotic fibrosarcoma) has been shown to increase the binding affinity with ARE, which leads to the transcription of phase II enzymes (Itoh et al., 1997). These enzymes play important roles in endogenous protection systems by neutralizing the harmful by-products of oxidative stress (Nguyen, Sherratt, & Pickett, 2003). A downstream gene in this pathway, Gsta3 (glutathione S-transferase alpha 3), was the most highly expressed gene in differentiating neural precursor cells following 25 μM quercetin treatment. In addition to Nrf2 upregulation, Keap1 (kelch-like ECH-associated protein 1) was also upregulated in the quercetin-treated group. This protein is the opponent of Nrf2 through facilitating the nucleo-cytoplasmic shuttling mechanism of Nrf2 and subsequently promoting its ubiquitinoylation and degradation (Nguyen, Sherratt, Nioi, Yang, & Pickett, 2005). Bell and colleagues demonstrated that the overexpression of Nrf2 inhibits neurite outgrowth and neuronal maturation (Bell et al., 2015). Therefore it is likely that the upregulation of Keap1 expression upon quercetin treatment, counteracts the detrimental effect of excessive Nrf2 accumulation.

The observation that quercetin up-regulated the intrinsic antioxidative pathways is interesting in this context. However, the involved pathways extend beyond the anti-oxidative effects that initially had been considered to be central to the biological effects of these compounds. The *in vivo* data are consistent with the *in vitro* results but still draw a complex picture. The reduction in proliferation, while survival and differentiation are promoted, point towards an interesting interaction deserving further studies.

Given the pro-neurogenic nature of quercetin, we sought to determine whether apple extracts display similar pro-neurogenic properties. We found that apple extracts from the flesh and to a lesser extent the peel increased neurosphere formation *in vitro*. Previous studies in aged animals found that apple supplementation prevented oxidative brain damage through increased endogenous antioxidative capacity, and also restored long-term potentiation and acetylcholine level, concurrent with improvement in behavioral tasks (Chan, Graves, & Shea, 2006; Tchantchou, Chan, Kifle, Ortiz, & Shea, 2005; Viggiano et al., 2006). A pilot study in human showed that consumption of apple juice improved behavioral symptoms in patients with Alzheimer’s disease (Remington, Chan, Lepore, Kotlya, & Shea, 2010). However, no significant cognitive change was observed after short term apple supplementation in healthy volunteers (Bondonno et al., 2014). We therefore investigated whether apple juice consumption could increase neurogenesis and enhance spatial learning and memory in mice. Despite observing pro-proliferative effects of apple extract *in vitro*, we found no effect of apple juice consumption on *in vivo* neurogenesis or spatial learning and memory performance. Given that the quercetin concentration in apple juice is very low (below 2 mg/L), resulting in an intake of approximately 0.6 mg/kg, we conclude that this is likely an insufficient concentration of active phytochemical to modulate neurogenesis with a measurable effect size. This however does not exclude a contributing effect to overall environmental effects on brain plasticity. Exposure to fresh and variable food is part of environmental richness, which has strong impact on the brain and its function.

Interestingly, however, we did identify additional previously unknown pro-neurogenic activity in the apple flesh, which is unrelated to flavonoids. Here we used an unbiased strategy to identify a candidate explaining part of that activity. In fraction IV of the 40 % MeOH fractionated apple flesh we identified DHBA as a pro-neurogenic factor. The 2,3-DHBA and 3,5-DHBA isoforms significantly increased the number of primary neurospheres, whereas the 3,4-DHBA isoform missed conventional statistical significance at *p* = 0.05. We found that the 3,5-DHBA isoform significantly increased cell survival in the adherent monolayer cultures under proliferation conditions and when added during the differentiation process in addition to increasing cell survival also significantly increased the neuronal lineage differentiation capacity. Hydroxybenzoic acid is an abundant compound in fruits, where it acts as a natural preservative (Aalto, Firman, & Rigler, 1953; Soni, Carabin, & Burdock, 2005). The food industry uses benzoic acid as preservative and in the European Union a declaration of its use must accompany a product. In contrast, naturally occurring and intrinsic DHBA has received relatively little attention as a potential signaling molecule. DHBA does, however, bind to the lactate receptor, Gpr81/Hcar1 (Cai et al., 2008), which, as we show, is present on neural precursor cells *in vivo* and *in vitro*, suggesting a direct mode of action. At the same time, given the role of lactate as both a signaling molecule and a source of energy for neurons, the discovery of an intrinsically active agonist to the receptor on neural stem cells is of further relevance (Lev-Vachnish et al., 2019; Wang et al., 2019). Given that it is an abundant dietary compound we also investigated whether 3,5-DHBA could also modulate *in vivo* precursor proliferation and neurogenesis. Here we found that 3,5-DHBA not only increased precursor cell proliferation and the number of newly born neurons, but also increased the maturation rate of these newborn cells. This appeared to be a concentration-dependent effect, with the most effective dose being 62.5 mg/kg and no effect observed at the highest concentration tested (250 mg/kg). In accordance with our results, another study also observed that administration of high concentrations of 3,5-DHBA (mg/kg) had no effect on the long-term survival of newly born neurons (BrdU^+^/NeuN^+^ cells approximately 7 weeks after the first dose of DHBA; (Lev-Vachnish et al., 2019). However, in that study lower concentrations of DHBA were not examined.

In both peel and flesh there will be more than one pro-neurogenic compound, our results are thus exemplary. Fruits contain numerous flavonoids and related substances and do so in relatively large quantities and varieties. This variety has especially struck researchers as interesting in that besides a shared biological activity across the range of molecules there might also be specific effects of individual natural compounds. These effects appear to more or less directly affect gene expression (for example via Sirt1 in the case of Resveratrol or PPARg in the case of EGCG). Our point is thus not that flavonoid activity would be absent from the flesh or that DHBA might not also be active in the peel. But neither compound alone, nor in combination, will explain the entire biological effect of exposure to food-born environmental signals. As can be seen from our *in vivo* experiments with quercetin versus those with apple juice, within the physiological ranges found in a fruit, the concentrations are too low to exert a unique (and thus, from an etiological perspective, presumably exaggerated) effect. But perhaps the neurogenic signals present in, for example, fruits might be maintenance signals and not elicit strong deviations from baseline. The *ex vivo* data, indicate that pro-neurogenic activity exists at low concentrations, but what remains to be elucidated is how the complex concert of many related factors achieve an effect. This will require a systems biology approach that goes beyond the scope of our present study.

The other critical question is, what the ethologically relevant function of this signaling might be? While there have been many studies on nutritional factors, including flavonoids, on brain health and plasticity, including a few preclinical studies on adult neurogenesis, the results regarding potential benefits, especially when translated to humans, have been mixed. However, the expectations towards such trials might have been exaggerated, in that natural compounds might be important environmental co-factors supporting an intrinsic homeostasis or baseline control rather than being strong extrinsic regulators above such baseline level. The mice had received a balanced, albeit constant diet. If we consider the observed effects as a means of communication between environment and the brain that results in adjusted plasticity, the exact situation in which individual such signals become measurable in their contribution to the complex picture remains to be identified. But our study suggests that such signaling exists. Our experiment with whole apple juice goes into that direction. In addition, our negative finding is valuable in that it confirms the absence of manifest non-specific effects (e.g. from sugar content).

We have shown that both flavonoids and DHBA are pro-neurogenic, not only by activating precursor cell proliferation but also through promoting cell cycle exit, cellular survival, and neuronal differentiation. One is tempted to speculate that the food-borne signal is part of a feedback loop between the environment and the individual foraging in it. Learning requires plasticity, but also stability, once relevant information has been gathered. Adult hippocampal neurogenesis provides flexibility in that it promotes the integration of novel information into pre-existing contexts. Searching for food has been a major task throughout evolution. It would be interesting, if via co-evolutionary mechanisms fruit would signal to the gatherer that their search has been successful and something has been learned.

## Materials and Methods

### Animals

For *in vitro* experiments, male and female C57BL/6JRj mice were purchased from Janvier and subsequently housed at the animal facility of Medizinisch Theoretisches Zentrum at Technische Universität Dresden, Germany. Animal handling in these experiments was conducted in accordance with the applicable European (86/609/EEC) and National (Tierschutzgesetz) regulations and approved by the local ethical committee. The mice were anesthetized with isoflurane prior to decapitation and brain isolation. *In vivo* experiments with apple juice supplementation and the water maze task were performed at the Centre for Regenerative Therapies Dresden, Technische Universität Dresden following the protocol that is approved by Animal Experiment Committee of Free State Saxony (DD24-5131/207/27). *In vivo* experiments to study the effect of quercetin was performed in the National Institute of Psychiatry Ramón de la Fuente Muñiz, Mexico City. The C57BL/6NHsd mice were purchased from Harlan Laboratories, Mexico City and Balb/C mice were obtained from the animal facility of the National Institute of Psychiatry "Ramón de la Fuente Muñiz". All institutional and legal regulations regarding to animal ethics and handling were followed in accordance with the national regulations (NOM-062-ZOO-1999) and had the approval of the Ethics Committee on Experimentation of the "Instituto Nacional de Psiquiatría Ramón de la Fuente Muñiz" (CEI/C/009/2013). Mice were housed in standard laboratory cages (4-5 mice per cage) under 12 h light/12 h dark cycles at a temperature of 23 ±1 °C. The light/dark cycle corresponded to the timing of lights on (Zeitgeber time 0; ZT0) at 0700 hours and to the timing of lights off (Zeitgeber time 12; ZT12) at 1900 hours, respectively. The animals had access to food and water *ad libitum*. Mice were acclimatized to their new environment until the age of 10 weeks.

### In vivo quercetin treatment

Chlordeoxyuridine (CldU) 42.5 mg/kg was injected at the beginning of the experiment and Iododeoxyuridine (IdU) 57.5 mg/kg was injected 2 h before the animals were perfused with 4% paraformaldehyde (PFA). Quercetin 50 mg/kg (Sigma) was prepared freshly in propylene glycol and administered daily through oral gavage (total volume 200 μl). The control group received the same volume of propylene glycol as vehicle control. Following brain processing, IdU^+^ and CldU^+^ cells were visualized with immunofluorescence staining and subsequently counted using a fluorescence microscope (Apotome, 40x objective) throughout the entire rostro-caudal axis of the granular cell layer.

### In vivo apple juice supplementation

Pinova apples were purchased from Obsthof Schlage Dresden-Pillnitz, harvested in autumn 2014. Eight liters of apple juice were obtained from 13 kg of fresh apples, then aliquoted and stored at −80°C. After a 1-week acclimatization period, the mice were housed in three groups of five that received juice, sugar water or normal drinking water. The sugar water or apple juice was replaced daily. Food and liquid intake were measured daily and body weight was measured weekly. A single injection of BrdU was given on the first day of the experiment and the mice were perfused one day after the completion of the water maze task.

### Isolation of adult hippocampal neural precursor cells

The isolation procedure of adult hippocampal neural precursor cells was adapted from Babu et al. (2011). Under isoflurane anesthesia, mice were killed by cervical dislocation, the brains collected and placed in ice-cold PBS. The dentate gyrus was dissected as previously described (Walker & Kempermann, 2014). The tissue was dissociated using Neural Dissociation Kit-P (Miltenyi Biotec) following the manufacturer’s instructions and the cells resuspended in 1 ml of growth medium (Neurobasal medium containing 2% B27 supplements, 1 x GlutaMAX, 2 μg/ml heparin, 50 units/ml Penicillin/Streptomycin, 20 ng/ml EGF and 20 ng/ml bFGF-2). The cells were then plated either into one well of a PDL- and laminin-coated 12-well plate and incubated in 37°C, 5% CO_2_ for monolayer culture or into an uncoated 96-well plate for neurosphere culture. The tested compounds/extracts were dissolved in water or DMSO, depending on the solubility. When DMSO was used as the vehicle, the final concentration in the culture medium did not exceed 0.1 % and an equivalent volume of DMSO was added to control conditions.

### Adherent monolayer cell culture

Five to six male and female C57BL/6JRj mice were used to establish a monolayer culture of precursor cells. After tissue dissociation the precursor cells were plated into a PDL-/laminin-coated plate and incubated at 37°C with 5% CO_2_. Every second day, half of the medium was replaced with fresh proliferation medium with the full concentration of growth factors (20 ng/ml bFGF and 20 ng/ml EGF). After reaching 80% confluency, the cells were passaged. The cells were washed quickly with PBS (without Mg^2+^ and Ca^2+^), detached by incubating with 500 μl Accutase for 3 min at 37°C. and washed in 10 ml PBS. The cell pellet was resuspended in 1 ml proliferation medium, and the cells counted using a hemocytometer and then replated at a density of 10,000 cells/cm^2^. The cells used for experiments did not exceed 15 passages.

### Neurosphere assay

Freshly isolated precursor cells from the dentate gyrus were resuspended in complete neurosphere medium to reach the approximate density of 2 hippocampi in 10 ml medium, and subsequently filtered through a 40 μm nylon mesh cell-strainer (Walker & Kempermann, 2014). After adding the test compounds or vehicle, cells were plated in a 96-well plate and incubated at 37°C, 5% CO_2_ for 12 days. Neurospheres with diameters ≥ 40 μm were counted using an inverted light microscope.

### Cell cycle analysis with propidium iodide staining

Cells were harvested and fixed with ice-cold 70% ethanol. Following two PBS washes, the cell pellet was treated with 50 μl RNase A (100 μg/ml). Subsequently, 400 μl propidium iodide solution (50 μg/ml solution) was added and the cells incubated for 1 h in the dark prior to flow cytometric analysis.

### 7-aminoactinomycin D (7-AAD) cell death assay

Cells were harvested into 15-ml falcon tubes and washed twice with PBS. The cell pellet was resuspended in 400 μl PBS and treated with 20 μl (1 μg) of 7-AAD solution. As the positive control, 0.2% Triton X (final concentration) was added to the cell solution. Flow cytometric analysis was performed following a 10 min incubation.

### In vitro BrdU labeling and immunostaining

BrdU labeling followed by immunofluorescence staining was used to measure the proliferation rate of neural precursor cells *in vitro*. Cells were plated on PDL/laminin-coated glass coverslips in 24-well plates with a seeding density 10,000 cells/cm^2^ and incubated in proliferation medium for 48 h. The medium was replaced with fresh proliferation medium containing test compounds or vehicle (DMSO) and further incubated for another 24 h. BrdU (10 μM) was applied to each well for 2 h prior to fixation. The cells were fixed with ice-cold PFA 4 % for 10 min and subsequently washed three times with PBS. Prior to BrdU immunofluorescence staining, the fixed cells on coverslips were washed twice with NaCl 0.9 % and then incubated in 1N HCl at 37°C for 30 min followed by one wash with 0.1M borate buffer, pH 8, and two washes with PBS. The blocking step was performed by incubating the cells in blocking solution (10% donkey serum and 0.2% Triton X-100 in PBS) at room temperature for 1 h. Subsequently, anti-BrdU antibody (1:500) in antibody solution (3 % donkey serum and 0.1 % Triton X-100) was dispensed on each coverslip and incubated either for 2 h at room temperature or overnight at 4°C. After three washes with PBS, the cells were incubated with fluorescent secondary antibody (1:500, in 3 % donkey serum and 0.1 % Triton X-100 for 90 min at room temperature. Following two washes with PBS cells were incubated for 10 min with Hoechst 33342 (1:3000 in PBS). After a final two washes with PBS, the coverslips were mounted on glass slides using Aqua-Poly/Mount for fluorescence microscopy. Cell counting was performed using a Leica DM5500B fluorescence microscope with StereoInvestigator (Microbrightfield) software. The number of BrdU-positive cells in a given population of cells was quantified from several visual fields and the average was presented as the cell percentage.

### Immunofluorescence staining of cells on coverslips

Standard protocol for staining cells on coverslips was performed by blocking the non-specific binding with blocking solution for 1 h at room temperature. Overnight incubation at 4°C with primary antibodies in PBS plus solution was performed immediately after removing the blocking solution, without washing step. After 3 times washing with PBS, the cells were incubated in fluorescent secondary antibody (1:500, in PBS-plus solution) for 90 minutes at room temperature followed by twice washing with PBS. Counter staining was performed by 10 min incubation with Hoechst 33342 (1:3000 in PBS) followed by another two washes with PBS. Finally, coverslips were mounted on glass slides using Aqua-Poly/Mount for fluorescence microscopy. In HCAR-1 staining (membrane antigen), 0.2% Triton X blocking solution and PBS plus solution was replaced with 0.2 % Saponin. Antigen retrieval step was not necessary for staining the monolayer NPCs on coverslips.

### Fluorescence and DAB immunostaining for brain tissue

Mice were deeply anesthetized by intraperitoneal injection of xylazine and ketamine and subsequently perfused with 0.9 % NaCl and PFA 4 %. The brains were collected and immersed in PFA 4% solution for 24 h and subsequently equilibrated in 30 % sucrose solution. Serial coronal cryosections (40 μm) were cut using a microtome (Leica) and stored in cryoprotection solution at −20°C. Every sixth section of each brain were pooled in one series for immunostaining. For immunofluorescence staining, brain sections were blocked in blocking solution for 1 h at room temperature prior to overnight incubation of primary antibodies in 4°C. After a thorough washing step with PBS, incubation with fluorescence secondary antibody was done for 3-4 h at room temperature. The tissues were subsequently mounted on glass coverslips with Aqua Polymount. In DAB staining, the sections were initially incubated with 0.6% H_2_O_2_ for 30 min to quench endogenous peroxidase in the tissue. After thorough washing with TBS, incubation with 1 h blocking solution followed by overnight incubation with primary antibodies was performed. The sections were incubated with biotinylated secondary antibodies for 4 h at room temperature followed by the incubation with avidin/biotinylated enzyme complex solution (ABC elite) for 1 h at room temperature. Thorough washing steps were performed between incubation steps. The chromogen 3,3’-diaminobenzidine was used as the enzyme substrate to develop a dark brown color for visualization of the labeled antigens. The sections were mounted on gelatin-coated glass slide, dehydrated in Neoclear® solution and then covered with glass coverslip in Neo-mount® solution before microscopic analysis. For HCAR-1 immunofluorescence staining, antigen retrieval process using citrate buffer was performed prior to standard immunostaining protocol (using Saponin 0.2% for cells permeabilization). The sections were washed with TBS and then incubated in 10 mM citrate buffer for 5 min in 95°C (water bath). The process was repeated twice with 15-20 min interval by leaving the sections in room temperature to cool down. The standard immunofluorescence protocol was subsequently followed.

### Confocal microscopy and image analysis

On the LSM 780, images were acquired using a Plan-Apochromat 20x/0.8 air objective. DAPI, GFP, Cy3 and Cy5 were excited using the laser lines 405 nm, 488 nm, 561 nm and 633 nm, respectively. For emission detection the following wavelength areas were used: DAPI: 415–450 nm, GFP: 499–534 nm, CY3: 588–623 nm and CY5: 649–690 nm. Images were processed offline using Fiji (National Institutes of Health) and Adobe Photoshop CS5®(Adobe Systems Incorporated). The image composites and the figures were assembled using Adobe Illustrator CS4.

### Cell quantification in brain sections (BrdU-, CldU-, DCX- or IdU-labeled cells)

The number of labeled cells was determined in series of every 6th section. Positive cells were counted exhaustively throughout rostro-caudal axis of the granule cell layer using 40x objectives.

The resulting numbers were multiplied by six to obtain the estimated total number of labeled cells per animal. For phenotyping, the number of colocalization between NeuN^+^ or S100β^+^ cells was determined in randomly selected 100 cells for each dentate gyrus and displayed as a cell percentage.

### Resazurin survival assay

Stock solution of 0.5 mM was prepared by dissolving Resazurin dye (7-hydroxy-3H-phenoxazin-3-one 10-oxide) in water and filter sterilizing. The final working concentration (0.05 mM) was prepared by diluting the 1-volume of stock solution in 9-volume of growth medium. The cells were incubated in this working solution for 2 hours in 37°C and the fluorescence intensity was subsequently measured in a plate reader at 560 nm Ex/590 nm Em filter settings. As the background, the solution was incubated in the wells without cells. After subtraction of all the fluorescence values with background value, the relative values were obtained by normalizing the fluorescence values of the treated group to the control group.

### Western Blot

For western blot analysis, the cells were collected in lysis buffer with protease inhibitor cocktail and disrupted by trituration using a syringe fitted with a 20G needle. The proteins were separated by SDS-PAGE gel electrophoresis and transferred to nitrocellulose membranes. The membranes were blocked and then probed with primary antibodies and HRP-labeled secondary antibodies. Protein bands were detected using a horseradish peroxidase/chemiluminescence system (ECL Western Blotting Substrate, Pierce) and visualized on Hyperfilm ECL (Amersham). The films were scanned and band intensities were determined by using ImageJ (http://imagej.nih.gov/ij/).

### RNA microarray

The total RNA from monolayer cell culture treated with quercetin was prepared using the Qiagen RNeasy mini kit, following the manufacturer’s protocol. The total RNA from primary cells treated with apple extract was prepared using Qiagen RNeasy micro kit, followed by amplification step using Ovation® Pico and PicoSL WTA Systems V2 and Encore® Biotin Module as described by the manufacturer. Prior to microarray analysis, the quality and integrity of RNA was assessed using a Bioanalyzer. RNA samples were amplified using the TotalPrep™ RNA Amplification Kit (Illumina) and hybridized to MouseWG-6 v2.0 Expression BeadChips (Illumina). Raw data were preprocessed with quantile normalization in R/Bioconductor using the package beadarray. The array was reannotated by querying each probe sequence against the mm9 mouse genome using the BLAT algorithm (kindly supplied by Dr. Jim Kent; http://hgwdev.cse.ucsc.edu/~kent/exe/linux). The genomic position of probes returning a single hit was then used to assign the probe to an NCBI EntrezGeneID. Probes targeting the same GeneID were collapsed as means to yield data for 21155 unique genes. Using the Benjamini-Hochberg method the gene expression result was corrected with the adjusted p value < 0.05 as the significance cut off and then visualized using a volcano plot. The gene expression was presented as “expression relative to control" which was the result of log2 fold change. The value “1” indicates 2 fold change of differentially upregulated expression and “-1” indicates 2 fold change of differentially downregulated expression. The list of differentially expressed genes was enriched into KEGG pathways (Kanehisa and Goto, 2000) and Wikipathways (Kelder et al., 2009) using WebGestalt, a web-based gene enrichment analysis tool. The microarray expression dataset used for analysis, as well as the raw data files, have been deposited in the GEO repository (http://www.ncbi.nlm.nih.gov/geo/) with the accession GSE150803.

### Real time PCR

Total RNA was prepared using the Qiagen RNeasy mini kit. The adherent cell cultures were washed once with PBS and then harvested using a cell scraper and RLT buffer. Subsequently, RNA was extracted following the manufacturer’s protocol, including the optional Dnase I step. The concentration and the purity of RNA were analyzed using a NanoDrop 1000 spectrophotometer. For reverse transcription, 1 μg of RNA was used. The cDNA was prepared using Superscript II kit using oligo(dT) primers following the manufacturer’s protocol. A cDNA negative reaction was also included without addition of reverse transcriptase enzyme to ensure no genomic DNA contamination. The quantitative real-time RT-PCR (qPCR) analysis for Bcl2l1 and Bcl2l11 was performed using a SYBR Green PCR Mix (Qiagen) following the manufacturer’s protocol. The template cDNA was prepared as above and each reaction contained cDNA from 0.1 μg total RNA. Thermal cycling and fluorescence detection were carried on an Opticon 2 DNA Engine (MJ Research). The quantification was done using the ΔΔCt method after normalizing to the β-actin housekeeping gene. The ΔΔCt shows the relative change of gene expression in the treated group compared to the control group.

### Apple extraction and fractionation

The extract was prepared from Pinova apple cultivar that was obtained from a local apple orchard (Obstgut Dreßler Sobrigau, Kreischa) (Schlage Obsthof) in autumn 2011. The peel was separated from the flesh using an apple peeler. The apple peel (100 g fresh weight) and flesh (200 g fresh weight) were separately mixed in 200 ml 80% methanol using a kitchen blender. The apple sludge was filtered through Whatman filter paper Grade 1. The process was repeated 3 times. The solution was then centrifuged for 10 minutes, 6000g at 4°C to remove the small particles from the extract. The supernatant was collected and dried in rotary evaporator at 40°C until reaching maximum dryness. The extracts were tested for the activity in neurosphere assay and then stored in −20°C for further fractionation steps.

The first fractionation steps were performed using the solvent fractionation method. The extracts were dissolved in ddH_2_O and 1 volume of ethyl acetate was added. Using a separation funnel, the water and ethyl acetate phase was separated and collected. The solvent was evaporated and the water fraction was further separated in the mixture of water and butanol (1:1). From these steps, 3 fractions were obtained each from apple peel and the solvent in all fractions was evaporated until maximum dryness. The active water fraction from the butanol separation was further fractionated by column separation using a Reverse phase C8 column (Vydac). The fraction was dissolved in 20% methanol solution before being injected into the column and then eluted with 20%-, 40%-, 60%- and 100% methanol consecutively for 30 minutes each. The elutes were then dried using centrifugal evaporator prior to being tested in the neurosphere assay. The active fraction (the one collected in 40% methanol) was further fractionated using Eurosphere C18 column using a gradient system with 1% aqueous acetic acid and 100 % acetonitrile (ACN) as the solvent. The gradient started with 80% ACN, then after 20 minutes decreased to 50% ACN, further decreased to 30% ACN within 20 minutes and finally within 20 minutes to 5%, followed by an equilibration step to the initial run condition. In this separation technique, 4 fractions were collected from different time ranges, i.e. minute 0-10, 10-20, 20-30 and 30-60, respectively. The solvent was evaporated in centrifugal evaporator prior to the neurosphere assay. The active fraction from this step was then analyzed using UPLC-MS system for structure elucidation.

### Compound elucidation by UPLC/ESI-QTOFMS

Chromatographic separations were performed at 40°C on an Acquity UPLC system (Waters) equipped with a HSS T3 column (100 Å~ 1.0 mm, particle size 1.8 μm, Waters) applying the following binary gradient at a flow rate of 150 μl min-1: 0-1 min, isocratic 95% A (water/formic acid, 99.9/0.1 v/v), 5% B (acetonitrile/formic acid, 99.9/0.1 v/v); 1-16 min, linear from 5 to 50% B; 16-18 min, isocratic 95% B; 18-20 min, isocratic 5% B. The injection volume was 2.6 μl (full loop injection). Eluting compounds were detected from 190-500 nm using a photodiode array detector (Waters) and from m/z 100-1000 using a MicrOTOF-Q hybrid quadrupole time-of-flight mass spectrometer (Bruker Daltonics) equipped with an Apollo II electrospray ion source in negative ion mode. The following instrument settings were applied: nebulizer gas, nitrogen, 1.6 bar; dry gas, nitrogen, 6 l/min, 190°C; capillary, +4000 V; end plate offset, −500 V; funnel 1 RF, 200 V; funnel 2 RF, 200 V; in-source CID energy, 0 V; hexapole RF, 100 V; quadrupole ion energy, 3 eV; collision gas, argon; collision energy, 10 eV; collision RF 200/400 V (timing 50/50); transfer time, 70 μs; pre pulse storage, 5 μs; pulsar frequency, 10 kHz; spectra rate, 3Hz. Mass spectra were acquired in centroid mode. Calibration of m/z was performed for individual raw data files on lithium formate cluster ions obtained by automatic infusion of 20 μl lithium hydroxide (10 mM) in the mixture of isopropanol/water/formic acid, 49.9/49.9/0.2 (v/v/v) respectively at the end of the gradient using a diverter valve. For the acquisition of collision-induced dissociation mass spectra, appropriate precursor ions were isolated using the first quadrupole (isolation width m/z 4) and fragmented inside the collision cell applying collision energies in the range of 15-30eV.

### Total flavonoid assay

Total flavonoids in apple cultivars were measured using the sodium borohydride/chloranil-based assay as described previously (He & Liu, 2008). Dried samples and quercetin standards were reconstituted in 1 ml THF/EtOH (1:1,v/v) in a glass test tube. 0.5 ml of 50 mM NaBH4 and 0.5 ml of 74.56 mM AlCl3 solution, were added to each tube followed by a 30 minute mixing step on an orbital shaker at room temperature. An additional 0.5 ml of NaBH4 solution was added with continuing shaking for a further 30 minutes at room temperature. Subsequently, 2 ml of 0.8 M of cold acetic acid solution was added, thoroughly mixed and then incubated in the dark for 15 minutes. The tubes were then heated to 100°C with shaking for 60 minutes after adding 1ml of 20 mM chloranil. The tubes were cooled down using tap water, and the final volume was brought to 4 ml with methanol. Then, 1 ml of 1052 mM vanillin was added into each tube and mixed thoroughly. Finally, 2 ml of concentrated HCl was added into each tube and kept in the dark for 15 minutes after a thorough mix. For detection, 200 μl of final reaction solution was added into each well of a 96-well plate and the absorbances were measured using a microplate reader at 490 nm.

### Morris water maze task

Three weeks after apple juice supplementation, mice were trained to locate a submerged escape platform in a circular pool (2 m diameter). Water was made opaque with non-toxic titanium dioxide and kept at a temperature of 19-20°C. Each mouse received 6 trials a day for 5 consecutive days. The position of the platform was changed on day 4 to the opposite quadrant (reversal). The starting position was changed every day and remained constant for all trials each day. Mice were allowed to search for the platform for up to 120 seconds. At the end of each trial, mice were guided to the platform and allowed to remain there for 15 seconds. The first 30 second time frame on first trial of day 4 was used as probe trial. Swim paths were recorded using Ethovision (Noldus) and further analyzed using Matlab (the Mathworks, USA).

### In vivo 3,5-DHBA treatment

Thirty-four female Balb/C mice were used for 3,5-DHBA administration. 3,5-DHBA was prepared freshly every day and dissolved in 5% ethanol in saline solution (in 0.9% NaCl) to administer at doses of 15.625, 31.25, 62.5, 125 and 250 mg/kg body weight per mouse. Then, 3,5-DHBA was administered once daily for 14 d via intraperitoneal (i.p.) injection. One hour prior to the first 3,5-DHBA administration, mice received two BrdU injections (50mg/kg, i.p.), each separated by one hour.

### Categorization of dendritic morphology of doublecortin-positive cells

Doublecortin cells were classified according to their morphological appearance, as described previously (Plumpe et al., 2006). Six categories were established based on the dendrite morphology: DCX-labelled cells; a) without dendrites; b) showing shorter dendrites than the soma size; c) with dendrites slightly larger than the soma size; d) with dendrites size longer than category "c"; e) with more mature appearance showing one primary dendrite with one branching point (node); and f) with mature appearance showing more than three nodes and reaching the molecular layer. Absolute quantification was performed by calculating the percentage of cells in each category and the total number of DCX-positive cells.

### Statistical analysis

Statistical analysis was performed using Graphpad Prism software (version 5.0, GraphPad Software Inc.). Depending on the number of groups, statistical analysis was performed using t-test, one-way or two-way ANOVA. One sample t-test was performed to analyze 2 groups, whereby each replicate in the treatment group is presented as normalized to control. Therefore the control group has only one value, which was then set as the hypothetical value. The Dunnett’s test was used as the post hoc test in one-way ANOVA. Microarray data analysis was performed using R for Mac OS X (3.1.1). The false discovery rate (FDR) method was applied to perform multiple comparison tests on the microarray data. All data are shown as means with standard error (SEM), unless stated otherwise.

## Acknowledgments

This project was funded by the Deutsche Akademischer Austauschdienst, Deutsche Forschungsgemeinschaft Sonderforschungsbereich 665, Bundesministerium für Bildung und Forschung, Consejo Nacional de Ciencia y Tecnologia and the Brazil Family Foundation for Neurology.

